# DNA microbeads for spatio-temporally controlled morphogen release within organoids

**DOI:** 10.1101/2024.01.10.575045

**Authors:** Cassian Afting, Tobias Walther, Joachim Wittbrodt, Kerstin Göpfrich

## Abstract

Organoids have proven to be powerful *in vitro* model systems that mimic features of the corresponding tissue *in vivo*. However, across tissue types and species, organoids still often fail to reach full maturity and function, because biochemical cues cannot be provided from within the organoid to guide their development. The establishment of such tools has been identified as a major goal of the field. Here, we introduce DNA microbeads as a novel tool for implementing spatio-temporally controlled morphogen gradients inside of organoids at any point in their life cycle. The DNA microbeads are formed in a simple one-pot process, they can be stored for a year and their viscoelastic behavior and surface modification is tunable to mimic the corresponding tissue. Employing medaka retinal organoids and early embryos, we show that DNA microbeads can be integrated into embryos and organoids by microinjection and erased in a non-invasive manner with light. Coupling a recombinant surrogate Wnt to the DNA microbeads we demonstrate the spatio-temporally controlled release of the morphogen from the microinjection site, which leads to the formation of retinal pigmented epithelium while maintaining neuroretinal ganglion cells. We were thus able to bioengineer retinal organoids to more closely mirror the cell type diversity of *in vivo* retinas. The DNA microbead technology can easily be adapted to other organoid applications for improved tissue mimicry.

## 1 Introduction

Organoids, derived from embryonic and (induced-) pluripotent stem cells, became a widely used tool in basic research, human disease modeling and personalized medicine since their emergence several decades ago. Organoids have been established for a variety of organs, among others, retina[1], brain[2], intestine[3], liver[4], pancreas[5], kidney[6], lung[7], thyroid gland[8], mammary glands[9] and prostate[10]. Retinal organoids (RO) specifically have been assembled and studied using embryonic and (induced-) pluripotent stem cells from mice, humans and fish. Among them, medaka (*Oryzias latipes*) fish RO develop by far the fastest, can be derived from easily producible transgenic reporter lines and studied in close comparison to the embryo of the corresponding developmental stage due to medaka’s transparent eggs.[1,11–12] Medaka RO are therefore particularly well suited for the development and implementation of new tissue engineering technologies.

While organoids, including RO, share many of their properties with their *in vivo* counterparts, end-point morphology, cell type diversity and functionality have proven difficult to replicate in their entirety as of now. Generally, culture conditions and the addition of extracellular matrix components can provide somewhat appropriate signaling that allow stem cells to undergo proliferation, differentiation and migration towards complex 3D architectures, but cues are regularly not spatially organized. This lack of spatial organization of morphogen gradients is one of the key factors limiting the organoids full emulation of the respective organ and keeping them from being a more physiologically relevant model system.[13] Using engineered materials for spatiotemporal delivery of bioactive cues to ultimately guide organoid development could be a promising avenue to properly address this limitation.[14–15]

Thus far, morphogen gradients have mainly been implemented in stem cell culture by microfluidic devices[16–19], patterning of hydrogels with biochemical cues[20–24] and integration of transgenic cellular signaling centers at an organoid’s pole[25]. Yet, these approaches can only provide unidirectional slopes of morphogen gradients from the outside to the inside of a respective organoid, always exposing the outer cell layers of an organoid to higher concentrations of morphogens than the inner cell mass. To create spatially discrete, organoid-internal morphogen sources and thus reversed gradients, the utility of micro-/nanoparticles in co-aggregation during early spheroid assembly has been explored previously.[26–29] Utilizing stem cell aggregate merging techniques, broad spatial control over microparticle mediated morphogen release in merged aggregates has been achieved.[29] Nevertheless, this technique gives the user no direct and precise spatial or temporal control, it is optimized for early organoid assembly and has only limited, if any, applicability in mid- to late-stage organoid culture. As such, better control over the onset of morphogen gradients as well as new and broadly applicable techniques for morphogen delivery are needed.

DNA hydrogel materials have gained much popularity in recent years owing to their simple programmability via sequence-specificity.[30–34] In this way, versatile DNA-based materials with controllable stiffness[35–39] and chemical modification e.g. by click chemistry[40–41] with pH-[40,42–43] or light-responsivity[44–48] have been created, including DNA droplets that form by liquid-liquid phase separation[40,49–56]. Pioneering studies show their utility as tools for the uptake and delivery of molecular cargo[49,57] even in living systems[40,50,58]. However, the potential of DNA microbeads as a tool for the engineering of organoids remains largely unexplored.

Here, we present DNA microbeads as a modifiable DNA hydrogel material that can be integrated as a spatially discrete and temporally controllable source of morphogen inside of an organoid at any point in its life cycle. The DNA microbeads are formed in a templated manner in a scalable one-pot reaction. Further, utilizing two orthogonal DNA nanostructures and a DNA linker allows us to incorporate multiple chemical modifications into the DNA microbeads at once. We thus achieve the formation of stable, cell-sized, stiffness-adaptable and chemically modifiable DNA microbeads, which can be delivered and integrated into the interior of early medaka embryos and medaka RO by microinjection.

Microinjected DNA microbeads do not influence the normal organoid development and are non-invasively erasable after tissue integration by light-triggered breakdown. To induce spatially organized morphogen gradients from within the organoid, the DNA microbeads are functionalized with a recombinant Wnt agonist as a morphogenic protein. By linking Wnt to the DNA microbeads via a photocleavable moiety, the morphogen is delivered and released within the organoid’s interior, ultimately creating morphogen activity gradients from the inside out in a temporally controlled manner.

This allows us to engineer retinal organoids more closely mirroring the cell type diversity of the *in vivo* retina, exemplifying how the presented tool can increase the complexity and phenotypic accuracy of organoid culture.

## 2 Results and Discussion

### 2.1 Templated formation of DNA microbeads allows for the tuning of material properties to match retinal organoid cells

To establish a generalizable tool capable of providing chemical cues from within the organoid, we set out to engineer DNA microbeads which fulfill several key requirements for the desired application.

**(i) Scalability.** The DNA microbead production should be simple and scalable, without the use of specialized equipment or expert knowledge, such that it can be performed in any laboratory with an interest in organoid culture. Long shelf life is an asset. **(ii) Tunable mechanics.** The mechanical properties of the DNA microbeads have to be tunable to mimic a diverse range of different tissues or to provide additional mechanical cues. **(iii) Microinjection-compatible.** The DNA microbeads have to be highly resistant to shear stress such that they can be microinjected into the organoid at any point in their life cycle. **(iv) Biocompatibility.** The microbeads should be stable in the organoids’ interior but also degradable on demand once they served their purpose in order to avoid undesired influence on organoid development. **(v) Chemically modifiable.** It has to be possible to attach multiple chemical cues onto the microbeads and release them on demand from the organoid’s interior in a non-invasive manner with full spatio-temporal control. This would enable user-defined morphogen gradients which guide development.

Hence, we first designed DNA microbeads which are compatible with our **Requirements (i-v)** and confirmed experimentally that the requirements are indeed fulfilled. We followed a DNA design consisting of three single strands, which, upon base pairing, form a branched, double-stranded DNA nanostructures with three arms, resembling the letter Y, hence termed Y-motif (**Figure 1A**).[59] These types of DNA nanostructures have been used as precursors for DNA hydrogels, as they can be interlinked via short sticky-end overhangs located at each end of the Y-motif arms.[38,48,60] We used two Y-motifs with orthogonal sticky-end overhangs. Upon addition of a single-stranded piece of DNA capable of binding to both sets of sticky-end sequences (DNA linker), these Y-motifs then bind to form a hydrogel network in bulk solution.[48,60]

**Figure 1.**
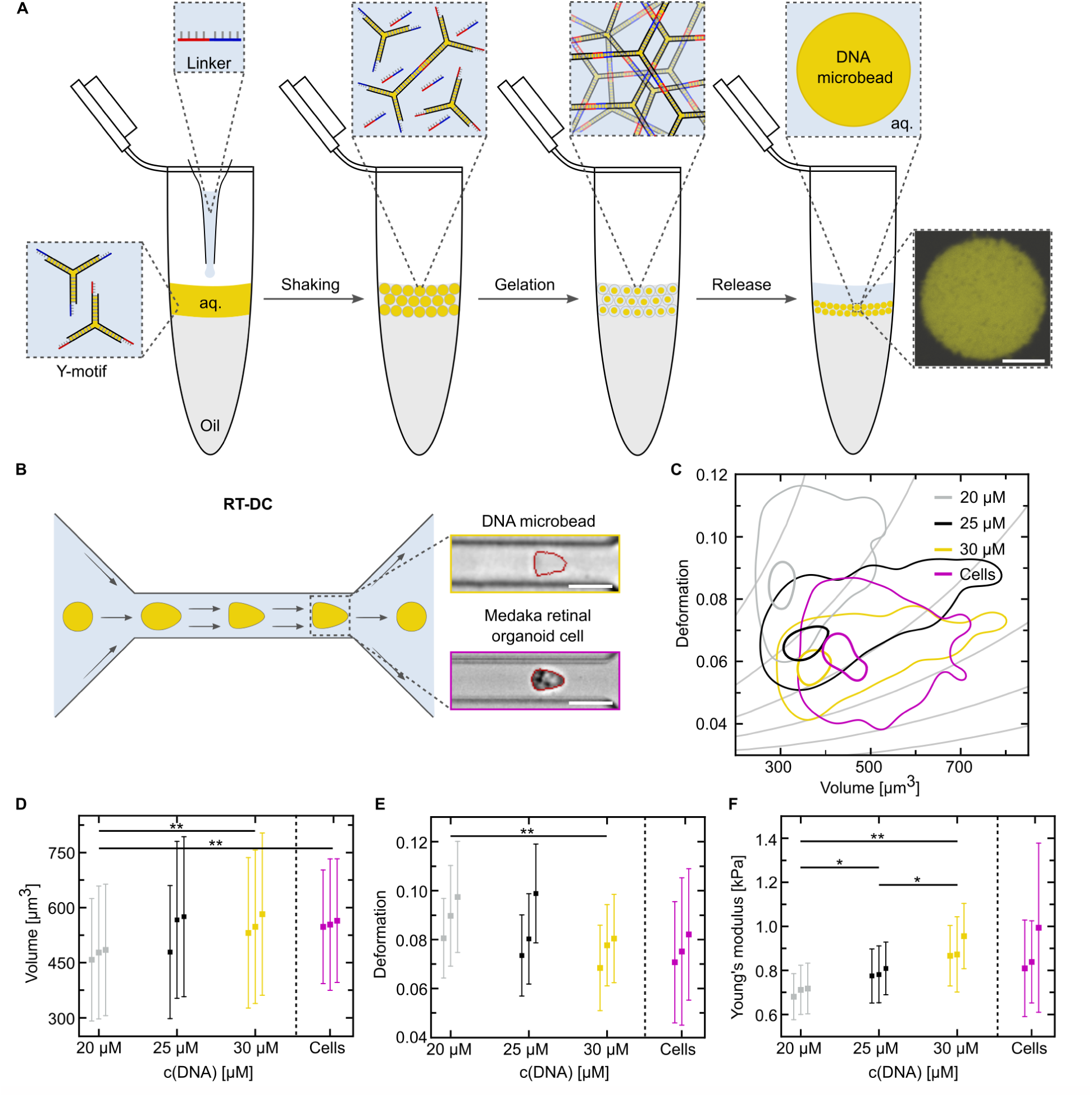
DNA microbead production and stiffness adaptation to RO cells. A) Schematic illustration of the DNA microbead production workflow. *Right:* Confocal microscopy image of a DNA microbead (λ_ex_ = 561 nm, Cy3-labeled DNA). Scale bar: 10 µm. B) Schematic illustration of RT-DC for high-throughput stiffness characterization. DNA microbeads or medaka RO cells are flushed through a microfluidic channel (width: 20 µm) and deform under the shear stress. Brightfield images showing a deformed DNA microbead and medaka RO cell, respectively. Scale bars: 20 µm. C) Deformation of DNA microbeads at various DNA concentrations (n_20µM_ = 32028, n_25µM_ = 41137, n_30µM_ = 38254) and medaka RO cells (n_Cells_ = 25853). The deformation of different populations of DNA microbeads and medaka RO cells is plotted over the corresponding volume using contour plots displaying the 50^th^ percentile (outer thinner line) and 95^th^ percentile (inner thicker line) of each measurement. D) Volume of the measured DNA microbeads and medaka RO cells. Statistically significant differences were detected for 20 µM and 30 µM DNA microbeads (**p-value: 0,002), and 20 µM DNA microbeads and medaka RO cells (**p-value: 0,001). E) Deformation of the measured DNA microbeads and medaka RO cells. Statistically significant differences were detected for 20 µM and 30 µM DNA microbeads (**p-value: 0,008). F) Apparent Young’s moduli of the measured DNA microbeads and medaka RO cells. Statistically significant differences were detected for 20 µM and 25 µM DNA microbeads (*p-value: 0,01), 20 µM and 30 µM DNA microbeads (**p-value: 0,001), 25 µM and 30 µM DNA microbeads (*p-value: 0,02) as well as for 20 µM DNA microbeads and medaka RO cells (*p-value: 0,035). Statistical significance was assessed using a linear mixed model. For each dataset (D-F), the mean values and corresponding standard deviations of three independent experiments are shown.

Based on this design, we first developed a scalable method for DNA microbead production (**Requirement (i)**). We realized that the encapsulation of the DNA strands into water-in-oil droplets allows for the droplet-templated formation of micron-sized DNA hydrogel beads. Since water-in-oil droplets can readily be formed by emulsification of an aqueous solution and an oil-surfactant mix, this method is highly scalable and does not require specialized equipment. The workflow, illustrated in **Figure 1A**, requires only minutes of manual labor. In brief, the aqueous solution containing the two orthogonal Y-motifs as well as the DNA linker is layered on top of an oil-surfactant (2 wt% perfluoropolyether–polyethylene glycol (PFPE–PEG) dissolved in perfluorinated oil) in a reaction tube. A droplet emulsion is created by shaking the tube. The DNA condenses into microbeads by self-assembly with the linker strands. In the final step, a destabilizing surfactant (perfluoro-1-octanol) is added to break up the emulsion and to release the ready-to-use DNA microbeads into the aqueous phase.[61] This facile formation stably produces large quantities of DNA microbeads in a one-pot reaction without specialized equipment or knowledge, making them an easy tool to implement in any laboratory (for details see Methods). We confirm that the DNA microbeads are stable after microcentrifugation and pelleting, which allows for facile buffer exchange. The microbeads form a stable, non-recovering network and can be stored in the fridge for at least one year (Figure S1). We could thus confirm that our method for the formation of DNA microbeads fulfills all aspects of **Requirement (i)**.

Note that the here presented formation method is new and considerably different from obtaining DNA microbeads by liquid-liquid phase separation which yield much more unstable condensates. In particular, DNA microbeads in the liquid state are unsuitable for our application inside of organoids.

In recent years, light has been shed on the importance of mechanical cues for cellular development, especially with regards to substrate stiffness and stem cell differentiation, highlighting the importance of **Requirement (ii)**, mechanical tuneability.[14,62–63] New tools for the engineering of living systems need to be tailored to accommodate physical parameters to mitigate unwanted effects or to provide mechanical cues on demand. As such, we set out to test whether the DNA microbeads can match the stiffness of organoid cells.

Real-time deformability cytometry (RT-DC) is used as a high-throughput microfluidic method to analyze differences in stiffness and response to fluid flow of varying cell types.[64] In RT-DC experiments, cells (and here DNA microbeads) are flushed through a microfluidic channel at kilohertz rates, causing them to deform due to viscous forces of the fluid flow (Figure 1B). The deformation is tracked in real time. Based on this deformation, the apparent Young’s modulus, i.e. the stiffness, of the measured cells or DNA microbeads can be deduced.[64–65] We thus utilized this method to analyze the behavior of DNA microbeads and medaka RO cells, in order to fine-tune the DNA microbeads to match the physical properties of the organoid cells.

For stiffness matching to the organoid cells, we assumed that the stiffness of the DNA microbeads can be tuned by varying the concentration of the Y-motif and the linker. We therefore prepared DNA microbeads at three different Y-motif concentrations (20 µM, 25 µM and 30 µM). RT-DC experiments revealed both an increase in overall DNA microbead volume as well as a decrease in measured deformation with higher DNA Y-motif concentration (Figure 1C, D, E).

While significant differences in volume were found between 20 µM and 30 µM DNA microbeads as well as between 20 µM DNA microbeads and medaka RO cells (Figure 1D), average overall volumes between the 25 µM and 30 µM DNA microbeads were comparable with those of medaka RO cells, with 30 µM DNA microbeads displaying an almost identical average overall volume (V_20µM_ = 473 ± 303 µm^3^, V_25µM_ = 540 ± 354 µm^3^, V_30µM_ = 553 ± 366 µm^3^, V_Cells_ = 555 ± 290 µm^3^).

For the deformation of the different DNA microbeads and medaka RO cells, we obtained a similar picture, with significant differences detected between the 20 µM and the 30 µM DNA microbeads. A clear trend was also visible from the overall averages of deformation, that the 30 µM DNA microbeads might be the closest fit to the medaka RO cells. As such, the 25 µM DNA microbeads deformed much more than the 30 µM DNA microbeads or medaka RO cells, which again showed almost identical overall average values (D_20µM_ = 0.089 ± 0.035, D_25µM_ = 0.084 ± 0.032, D_30µM_ = 0.075 ± 0.03, D_Cells_ = 0.076 ± 0.047).

This trend became clearer when also analyzing the apparent Young’s moduli of the different DNA microbeads and cell samples. Here, significant differences were detected between all three of the tested DNA Y-motif concentrations, while no significant difference in apparent stiffness was detected between the 30 µM DNA microbeads and the medaka RO cells. Moreover, the overall average stiffness value for these two samples was again almost identical still (YM_20µM_ = 0.7 ± 0.19 kPa, YM_25µM_ = 0.79 ± 0.26 kPa, YM_30 µM_ = 0.89 ± 0.26 kPa, YM_Cells_ = 0.88 ± 0.48 kPa). All measured values for the volume, deformation and Young’s moduli of all samples are detailed in Tables S1 - S3.

With dynamic mechanical analysis measurements by microindentation of the DNA microbeads, we further found that they exhibit a frequency-dependent strain response similar to cells[66] (Figure S2). The DNA microbeads thus also mimic the viscoelastic solid behavior and distinct frequency-response of soft tissue cells making them an ideal cell mimic.[67–68]

Note that by changing the arm length of the Y-motif and by utilizing DNA tiles to create DNA microbeads, we were able to extend the achievable range of stiffnesses beyond the range relevant for RO - ranging from 20 Pa, mimicking the stiffness of cortical neuronal cells[69], up to 2 kPa, more closely resembling mesenchymal stem cells or fibroblasts[70–71] (Figure S2).

We thus confirmed that the DNA microbeads exhibit mechanical tunability fulfilling **Requirement (ii).** Due to the excellent match of the mechanical properties of the organoid cells, we selected the 30 µM DNA microbeads to be used in all further experiments. Note that the stiffness and viscoelastic properties of DNA microbeads had previously not been characterized. It is thus remarkable that they mimic the mechanical properties of cells so closely, making them ideally suited for our envisioned use case in organoid culture.

### 2.2 DNA microbeads can be delivered and integrated into *in vitro* and *in vivo* medaka retinas

Having demonstrated that DNA microbeads can be produced in a scalable manner and have suitable mechanical properties, we test whether they are sufficiently stable for microinjection *in vivo* and *in vitro* (**Requirement (iii)**).

For this, we turned to the fast-developing vertebrate model, medaka fish (*Oryzias latipes*). For *in vitro* and *in vivo* DNA microbead integration, we developed experimental pipelines for solid microparticle microinjection into medaka RO and medaka embryos (see Methods for details). Similar as described previously[11], medaka RO were generated as a 4:1 mix of primary pluripotent embryonic stem cells from wildtype and transgenic (Atoh7::EGFP) embryos. Therefore, only a fraction of retinal ganglion cells was being reported for, making it easier to spot qualitative differences in cell numbers and distribution within individual organoids. In this way, the labeled retinal ganglion cells were used as a proxy for the overall formation of neuroretina in the organoids (hereafter referred to as neuroretinal ganglion cells). RO were microinjected with DNA microbeads at late day 1, shortly after Matrigel-induced onset of neuroepithelium formation (**Figure 2A**). Light sheet microscopy of plasma membrane stained (CellMask^TM^ Deep Red) organoids showed that the DNA microbeads integrated seamlessly into the organoid’s tissue environment (Figure 2B). Note that the DNA microbeads withstand the strong shear forces during microinjection without disintegration and are stable within the organoid system even under changing culture media and conditions (**Requirement (iv)**). Likewise, microinjected DNA microbeads were integrated into the corresponding developmental stage of the *in vivo* embryonic medaka retina (embryo stage 20: optic vesicle formation[72]; Figure 2C). Culturing microinjected organoids until differentiation onset at day 4 showed that the DNA microbeads were stable within the organoids (Figure 2D), while they were naturally broken down within the developing retina of the medaka embryo over the course of 6-9 h (Figure S3A). This neither affected the survival nor the gross morphological development of the embryos to hatchling stage compared to non-injected control embryos (embryo stage 40[72]; Figure S3). For RO, we whole-mount antibody stained DNA microbead microinjected RO at day 4 with common molecular markers for differentiated retinal cell type identities (Atoh7::EGFP [neuroretinal ganglion cells], Otx2 [bipolar cells and photoreceptors] and HuC/D [amacrine and neuroretinal ganglion cells]) and imaged them via confocal microscopy. Differentiated retinal cell type composition and patterning did not differ from the respective PBS-microinjected and non-injected controls in all of 25 organoids imaged across 3 independent experimental replicates (representative images in Figure 2D). Thus, neither the presence and integration of DNA microbeads within nor the microinjection into RO seemed to affect their normal development (**Requirement (iv)**).

**Figure 2.**
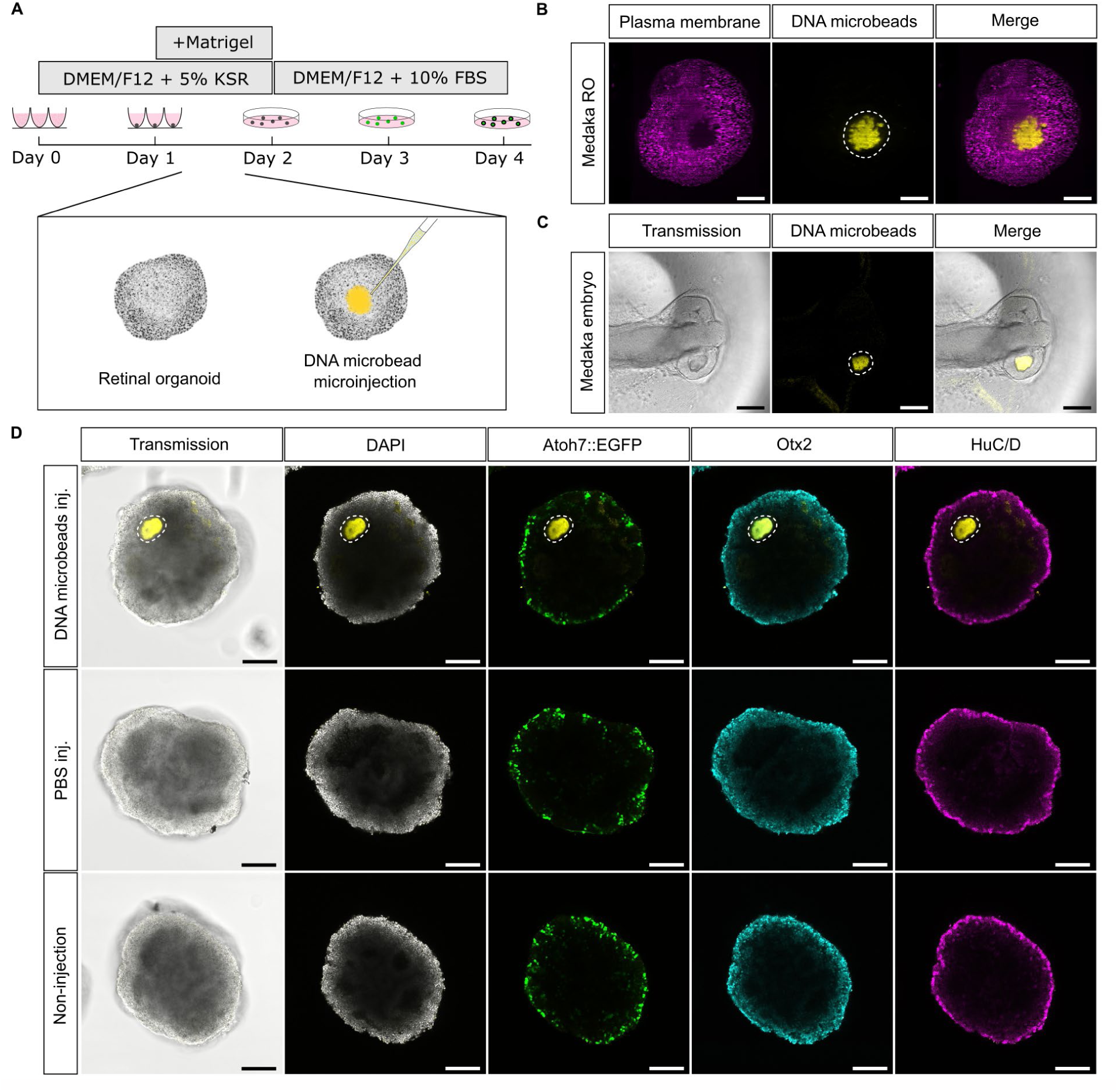
DNA microbead delivery and integration into the developing *in vitro* and *in vivo* medaka retina. A) Schematic illustration of medaka RO generation and time point of DNA microbead microinjection. B) Light sheet fluorescence microscopic image of a plasma membrane-stained day 3 RO post microinjection with DNA microbeads. C) Transmission image of an alive stage 20 medaka embryo 2 h after DNA microbead microinjection into its developing retina. D) Representative confocal images of whole-mount antibody stained day 4 RO (DAPI [nuclei], Atoh7::EGFP [neuroretinal ganglion cells], Otx2 [bipolar cells and photoreceptors] and HuC/D [amacrine and neuroretinal ganglion cells]) after microinjection with DNA microbeads (DNA microbeads inj.), PBS (PBS inj.) or being left uninjected (non-injection). Dashed white lines outline the DNA microbeads’ positions. Representative images from n = 25 organoids across 3 independent experiments. Scale bars: 100 µm.

### 2.3 DNA microbeads modified with a photocleavable group allow for non-invasive removal after tissue integration

Having confirmed that DNA microbeads are stable inside medaka RO and do not hinder their development, we next incorporated functionality into the DNA microbeads as a means of controlling their behavior inside the organoids. In particular, it is important that the DNA microbeads are not only stable as long as needed, they also have to be removable in a non-invasive way once they are no longer needed (**Requirement (iv)**). Adding a photocleavable (PC) moiety to the center of the DNA linker[48] allowed for the near-instantaneous breakdown of the DNA microbeads upon irradiation with 405 nm light with spatio-temporal control (PC-modified DNA microbeads; **Figure 3A/B**). The microbeads break down only in the illuminated area, while neighboring microbeads remain intact (Figure S4). We quantified the breakdown of DNA microbeads in solution following 60 s irradiation with a 405 nm laser, showing full degradation of the DNA microbeads within the 60 s (Figure 3B).

**Figure 3.**
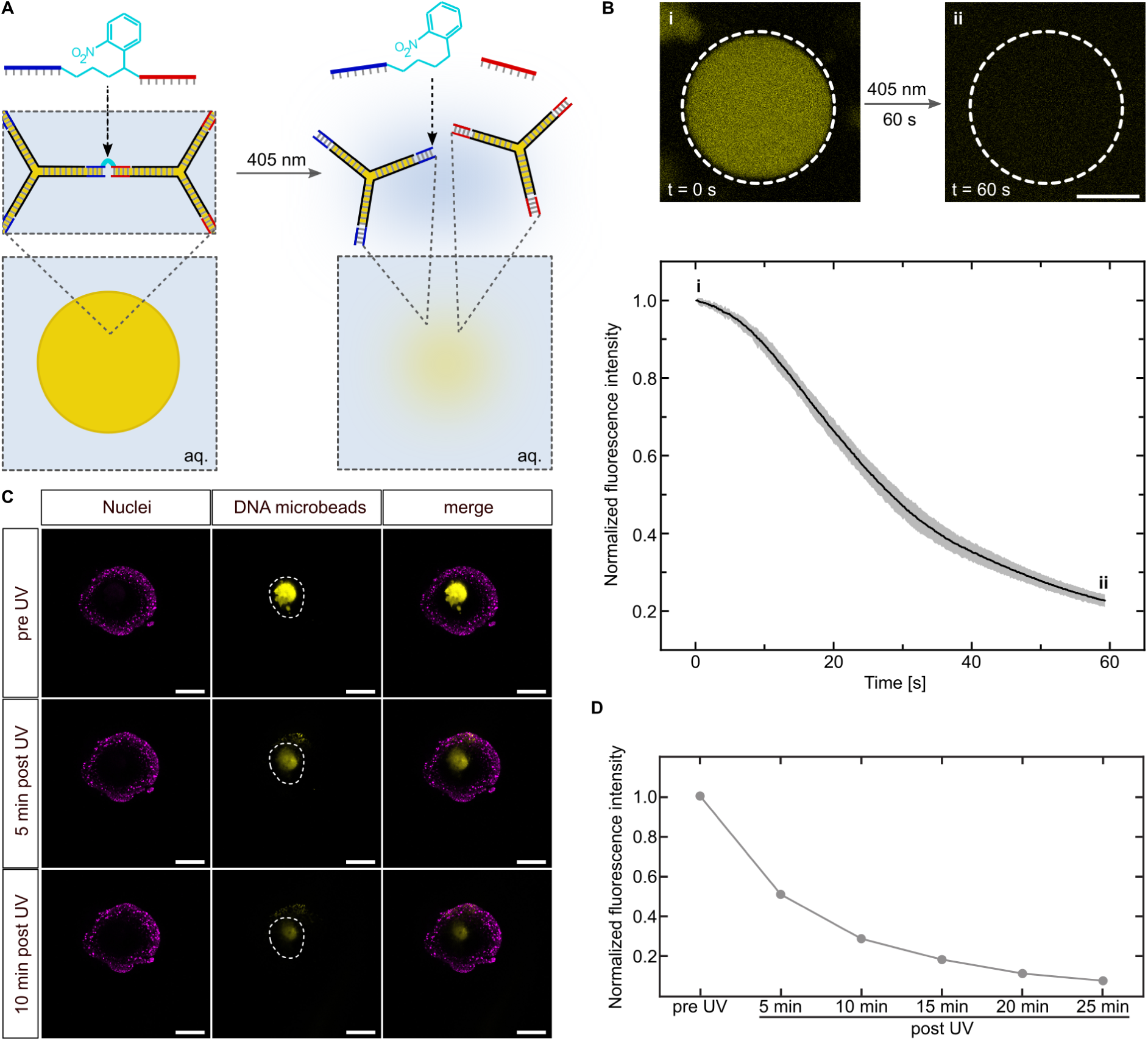
Photocleavable modification allows for the non-invasive removal of DNA microbeads from RO after their integration. A) Schematic illustration of the DNA microbead design with an internal photocleavable group (PC) in the DNA linker sequence. B) *Top:* Representative confocal images (λ_ex_ = 561 nm, Cy3-labeled DNA) of a PC-modified DNA microbead before (i) and after (ii) 60 s illumination with a 405 nm confocal laser (0.005 mW power). The white dashed line indicates the illuminated area. Scale bar: 20 µm. *Bottom:* Normalized fluorescence signal (mean ± standard deviation, 3 independent replicates analyzing 5 DNA microbeads each) plotted over the exposure time of 60 s. C) Representative confocal images of live nuclei stained, PC-modified DNA microbead microinjected RO before (pre UV), 5 min after (5 min post UV) and 10 min after (10 min post UV) exposure to UV light. Dashed white lines outline the DNA microbeads’ positions. Scale bars: 100 µm. D) Normalized fluorescence intensity of the DNA microbeads shown in C was plotted in 5 min steps over 25 min post UV exposure.

To test the light-triggered breakdown of PC-DNA microbeads in RO, we first stained the organoid’s nuclei with live nuclei stain (DRAQ5^TM^) and microinjected the PC-DNA microbeads, followed by timelapse confocal microscopy. Organoids were imaged prior to and after exposure to a short 1 min UV light pulse in 5 min increments. Upon UV light irradiation, the fluorescent signal of the DNA microbeads started to diffuse throughout the organoids ex- and interior and was gradually reduced to below the detection threshold within roughly 25 min (Figure 3C/D). Therefore, using the PC modification on DNA microbeads allows for their non-invasive removal after tissue integration with full user control. Whole-mount antibody staining, and confocal microscopy did not show any difference in retinal cell type composition and patterning of day 4 RO after PC-DNA microbead breakdown compared to UV light-treated controls. This was observed in all of 25 organoids imaged across 3 independent experimental replicates (representative images in Figure S5). Accordingly, neither the UV light regime nor the release of free DNA-motifs negatively affected normal medaka RO development. We thus gained control over the incorporation and breakdown of the DNA microbeads inside medaka RO, fulfilling also **Requirement (iv)**.

### 2.4 Photocleavable DNA microbeads permit a temporally and spatially controlled morphogen release within organoids

Finally, we demonstrated the utility of the DNA microbead system to guide organoid development by the targeted release of chemical cues (**Requirement (v)**). In particular, we aimed at releasing a Wnt agonist as a morphogen from the DNA microbeads within the RO. Canonical Wnt signaling has been described as a key factor in retinal pigmented epithelium (RPE) fate specification in early eye development.[73–75] Wnt agonists are frequently used in RO cell culture and are known to induce the formation of RPE which is otherwise occurring rarely and insufficiently. Currently, this comes with the unfortunate suppression of neuroretinal tissue[1] and therefore does not permit the full emulation of the *in vivo* retinal cell type diversity in RO cell culture across species.

Here, we used the extracellularly binding, next generation surrogate Wnt[76] (Wnt-surrogate), as a cell culture medium supplement in increasing concentrations. In accordance with literature on other agonistic Wnt molecules, supplementing increasing concentrations of Wnt-surrogate to the medium at day 1 of RO culture resulted in increasing amounts of RPE in the organoids, while at the same time heavily suppressing neuroretinal differentiation as visualized by the occurrence of neuroretinal ganglion cells (**Figure 4A/B**). Using bio-orthogonal DBCO-azide click chemistry[41,77–78], we then designed DNA microbeads with Wnt-surrogate proteins covalently attached via a photocleavable group to the DNA linker. We confirmed the successful linkage with sodium dodecyl sulfate polyacrylamide gel electrophoresis (Figure S6). This allowed for the incorporation of Wnt-surrogate into the microbeads, as modified DNA is readily taken up and incorporated into the DNA microbeads after formation (Figure S7). Wnt-surrogate could therefore be cleaved from the DNA microbeads upon irradiation with UV light, leaving the DNA microbeads intact (Wnt-DNA microbeads; Figure 4C, Figure S8). With microinjection of Wnt-DNA microbeads into and subsequent release of the Wnt-surrogate within the RO, we were then able to induce RPE formation, while neuroretinal ganglion cells were not notably suppressed in occurrence compared to control organoids without Wnt-surrogate release from the Wnt-DNA microbeads (Figure 4E, additional representative images in Figure S9, quantification of neuroretinal ganglion cell numbers in Figure S10). DNA microbeads thus allow us to provide morphogen gradients from within the organoid, which allow for more complex and realistic organoid development.

**Figure 4.**
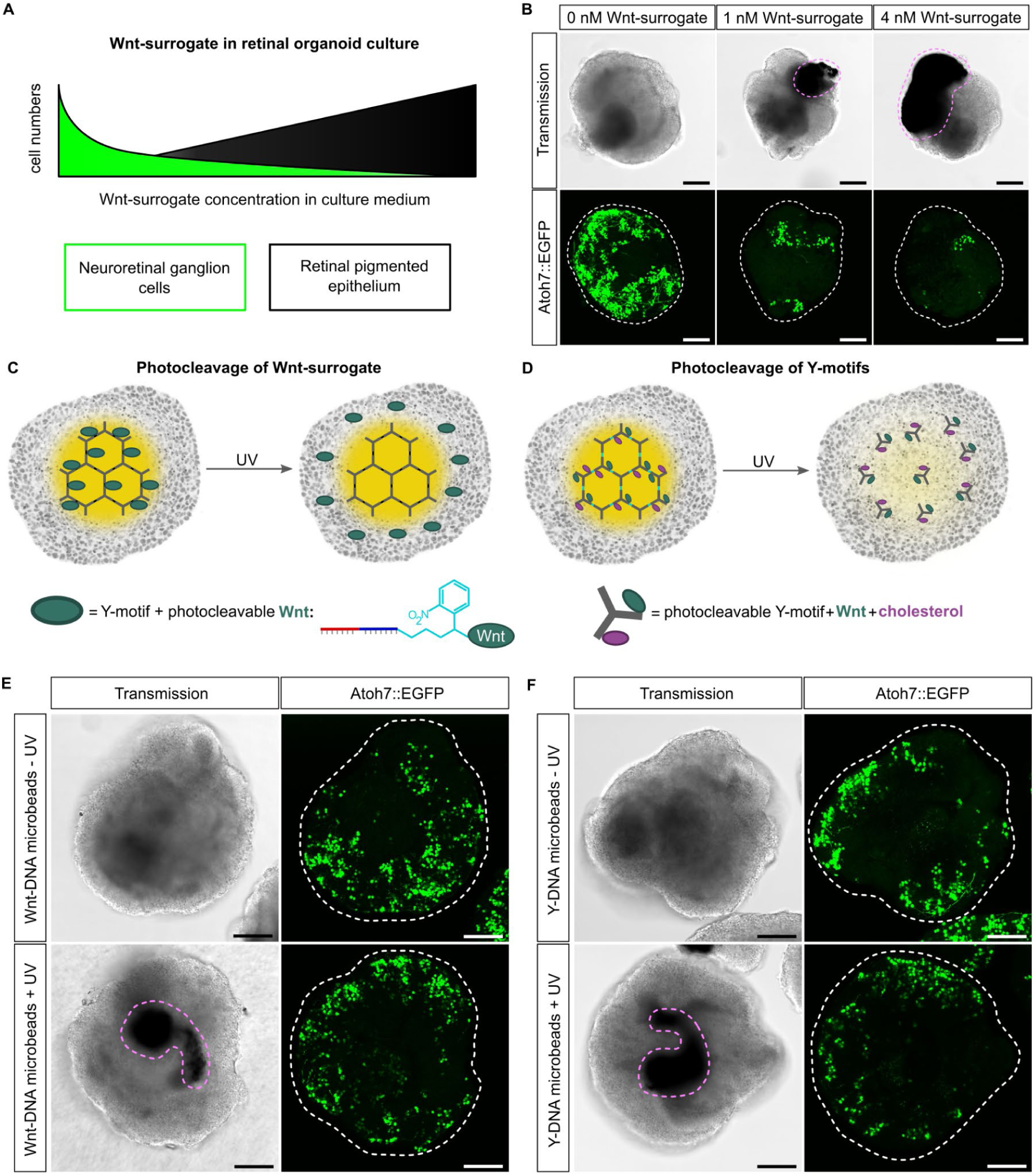
The photocleavable modification of DNA microbeads permits temporally and spatially controlled morphogen release within RO. A) Schematic illustration of retinal pigmented epithelium and neuroretinal ganglion cell numbers in RO with increasing concentrations of Wnt-surrogate in the culture medium. B) Representative confocal transmission and maximum intensity z-projection (Atoh7::EGFP; 35 slices at 5 µm distance) images of day 4 RO (n = 6 each) treated with 0, 1 and 4 nM Wnt-surrogate in the culture medium. C) Schematic illustration of the DNA microbead design with photoinducible Wnt-surrogate release from intact DNA microbeads. D) Schematic illustration of DNA microbead design with photocleavable Wnt-surrogate/cholesterol-Y-DNA-motif. Following UV-exposure, the DNA microbeads are broken down to release the Wnt-surrogate attached to cholesterol-modified DNA Y-motifs. E) Representative confocal transmission and maximum intensity z-projection (Atoh7::EGFP; 15 slices at 10 µm distance) images of day 4 RO after Wnt-DNA microbead microinjection and Wnt-surrogate release at day 1 (DNA microbead design illustrated in *C*). F) Representative confocal transmission and maximum intensity z-projection (Atoh7::EGFP; 12 slices at 10 µm distance) images of day 4 RO after Y-DNA microbead microinjection and Wnt-surrogate/cholesterol-Y-DNA-motif release at day 1 (DNA microbead design illustrated in *D*). Magenta dashed lines indicate retinal pigmented epithelium, while white dashed lines outline the shape of the respective RO. Representative images from n = 50 organoids (E, F) across 3 independent experiments. Scale bars: 100 µm.

Showcasing the possibility to add multiple functional moieties, we next added a cholesterol moiety to the DNA microbeads to enhance their anchoring on the cellular membrane. The Wnt-DNA microbead design was changed to where the Wnt-surrogate is released after DNA microbead breakdown while attached to the DNA-Y-motif and a cholesterol-moiety (Y-DNA microbeads; Figure 4D), to reduce its diffusivity. In this way, it is possible to release Wnt-surrogate and to remove the DNA microbeads in a single step. This design variant resulted in the same outcome in the RO (Figure 4F, additional representative images in Figure S9). Combining non-invasive removal of the DNA microbeads with the induction of organoid-internal Wnt-surrogate gradients highlights that a dual cargo release of different types of macromolecules is feasible with the DNA microbead technology, fulfilling also the final requirement (**Requirement (v)**). Off note, Wnt-DNA microbead microinjected organoids sometimes developed tiny hubs of RPE without UV light triggered Wnt-surrogate release. This might be due to Wnt-surrogate acting upon the cells directly adjacent to the Wnt-DNA microbeads even without release, although a minor non-detectable DNA microbead degradation cannot be ruled out as a cause.

In summary, we were efficiently inducing retinal pigmented epithelium on demand and adjacent to neuroretinal tissue in medaka RO. Strikingly and in contrast to systemic Wnt-surrogate treatment, the neuroretina was being effectively preserved. This is best explained by a Wnt-surrogate activity gradient emanating from the position of the microinjected DNA microbeads upon UV light exposure. Therefore, the exposure of the neuroretinal cells residing near the rim of the RO to the Wnt agonist was limited, differing from the standard approach of adding Wnt agonists to the culture medium and thus exposing the outer cells most. Our DNA microbead technology consequently enables the bioengineering of RO with a cell type composition more closely mimicking the *in vivo* retina, which otherwise cannot be achieved consistently and on demand via common culturing methods and interventions.

## 3 Conclusion

In this work, we show that DNA Y-motifs can be fabricated into stable, spherical microbeads by templated formation in water-in-oil droplets in a simple one-pot reaction without specialized equipment. The DNA microbeads can be fine-tuned to match the stiffness of a given cell type by varying their component concentrations. Stiffness-matched DNA microbeads are delivered and integrated into the interior of medaka RO by microinjection, without perturbation of the organoid’s expected differentiation and development. Modification of the DNA microbead structure with photocleavable groups permits the biocompatible UV light-triggered breakdown of assembled DNA microbeads even after integration into organoids. Functionalization of the DNA microbeads with a morphogen cargo using click-chemistry enables UV light-triggered cargo release from within the organoid’s interior. With that, we increase the cell type diversity of RO to match the cell type diversity of *in vivo* retinas more closely, which is not possible by supplementing the morphogen to the culture medium and thus creating a morphogen activity gradient with a slope from the outside in instead of vice versa.

Combining the possibility of cell-sized, stiffness-adapted DNA microbead microinjection with a UV light-triggered cargo release mechanism, this technology allows for unprecedented spatial and temporal user control in the bioengineering of organoids with internally provided morphogens throughout their life cycle. While this work presents a first proof-of-principle application of this mechanism using proteins, delivery of any click-chemically addressable molecule into tissues is feasible based on the same principle. The presented technology addresses the need for implementation of morphogen sources into 3D organoid cell culture of any developmental stage and more than ever before opens up their intricate interior microarchitecture to precise bioengineering efforts.

## 4 Experimental section

### Design and handling of DNA sequences

The sequences used to prepare the DNA Y-motifs YA and YB as well as the DNA linker were adapted from previous publications.[48,60] Binding of the different DNA strands was verified using NuPack.[79] DNA strands were purchased either from Integrated DNA Technologies (unmodified DNA, purification: standard desalting) or Biomers (modified DNA, purification: HPLC). All DNA, apart from fluorophore-labeled strands, was diluted in 10 mM Tris (pH 8) and 1 mM EDTA (Sigma Life Science) to yield 800 μM stock solutions. Fluorophore-labeled strands were diluted in MilliQ water to yield 800 μM stock solutions. All utilized DNA sequences are listed in Table S4, Supporting Information. The DNA stock solutions were stored at −20°C.

### Preparation of Y-motif DNA

The DNA Y-motifs (YA, YB) needed to form the DNA microbeads were produced via thermal-annealing of the three respective single-stranded DNA strands YA-1, YA-2 and YA-3 for YA, or YB-1, YB-2 and YB-3 for YB. The strands were mixed at equimolar ratios to yield a final concentration of the resulting Y-motifs of 150 µM. In all experiments, 4 mol% of Cyanine-3 (Cy3)-labeled YB-1 strand was added to the YB mixture in order to allow for fluorescence microscopy of the resulting DNA microbeads. The Y-motifs were annealed in a solution containing 1x phosphate-buffered saline (PBS, Gibco). Annealing was conducted in a thermal cycler (BioRad) by heating the samples to 85°C for 3 min and subsequently cooling the sample to 20°C using an increment rate of -0.1°C/s.

### Formation of DNA microbeads

DNA microbeads were created in a templated manner after encapsulation of the gelation solution into water-in-oil droplets. To form the DNA microbeads, the annealed Y-motifs YA and YB were mixed at equimolar ratios (20, 25 or 30 µM) in a solution containing 1x PBS. The DNA linker strand was then added to the solution in 3x excess to the Y-motifs. Immediately after the addition of the DNA linker, the mixture was added on top of an oil-phase containing 2 wt% perfluoropolyether–polyethylene glycol (PFPE–PEG, RAN Biotechnologies) dissolved in HFE-7500 (Iolitex Ionic Liquids Technologies) at a ratio of 1:3 aqueous-phase to oil-phase (e.g. 50 µL aqueous solution and 150 µL oil-mixture) and the reaction tube with the mixture flicked with a finger 8x to create an emulsion. The resulting water-in-oil droplet emulsion was left at 22°C room temperature for 72 h to ensure full gelation of the DNA microbeads. After this, the DNA microbeads were released by breaking the water-in-oil emulsion. To release the microbeads, a 1x PBS-solution was added on top of the droplet emulsion. Subsequently, the emulsion was destabilized by adding the surfactant 1H,1H,2H,2H-Perfluoro-1-octanol (Merck) on top of the buffer. This mix was left to incubate for 30 min before the resulting aqueous phase containing the DNA microbeads was taken off and transferred to a separate reaction tube. The DNA microbeads were stored at 5°C prior to their use and prepared fresh for each experiment.

### Confocal fluorescence microscopy of DNA microbeads

For imaging of the DNA microbeads, an LSM 900 confocal fluorescence microscope (Carl Zeiss AG) was used. For each experiment, the pinhole size was set to one Airy unit and a Plan-Apochromat 20x/0.8 Air M27 objective was utilized. All imaging was conducted at 22°C room temperature. For imaging, the DNA microbeads were deposited into custom-built observation chambers made from glass slides (Carl Roth) attached via double-sided sticky tape (Tesa) and sealed using two-component glue (twin-sil, Picodent). Prior to assembly of the observation chamber, the glass slides were coated for 5 min with poly(vinyl-alcohol) (50 mg/mL, Sigma Aldrich).

### Real-time deformability cytometry

Real-time deformability cytometry (RT-DC) was performed using an AcCellerator (Zellmechanik Dresden) mounted on an inverted AxioObserver microscope (Carl Zeiss AG) equipped with a 20x/0.4 Ph2 Plan-NeoFluar objective (Carl Zeiss AG). Images were acquired using a high-speed CMOS-camera (MC1362, Microtron).

To measure the DNA microbeads, a suspension of microbeads (100 µL) was strained through a 20 µm EASYstrainer filter (Greiner Bio-One) and pelleted in a reaction tube by spinning them down for 2 min with a C1008-GE myFUGE mini centrifuge (Benchmark Scientific). 80 µL of the supernatant were then taken off and discarded, and the remaining pellet of DNA microbeads was resuspended in 150 µL of CellCarrierB (Zellmechanik Dresden). The resuspended microbeads were then aspirated into a 1 mL glass syringe with PEEK tubing connector and PTFE plunger (SETonic) mounted on a syringe pump system (NemeSys, Cetoni). The DNA microbead-CellCarrierB solution was then injected into a Flic20 microfluidic chip (Zellmechanik Dresden) using PTFE-tubing (S1810-12, Bola). Through a second 1 mL glass syringe, CellCarrierB was injected into the Flic20 microfluidic chip as sheath flow for the RT-DC experiment. For all samples, two sequential measurements at 0.04 µL/s and 0.08 µL/s total flow rates (ratio of sheath-to-sample flow: 3:1) were run for a duration of at least 900 s each. The measurement software ShapeIn (version 2.2.2.4, Zellmechanik Dresden) was used to detect the DNA microbeads in real time. The pixel-size was adjusted to 0.68 µm/px, fitting the utilized 20x/0.4 Ph2 objective and all DNA microbeads imaged at the rear part of the flow channel ensuring regular deformation of each microbead. For each condition, triplicates were measured.

The same workflow was applied to dissociated medaka RO cells. In preparation for RT-DC, medaka RO were cultivated as described in “Generation of medaka derived retinal organoids” until late day 1. 48 organoids per experiment were then pooled into 2 ml tubes and washed multiple times with 1x PBS. Dissociation was performed by incubation in dissociation solution (1:1 dilution of 2.5% Trypsin (Gibco^TM^, Cat#: 15090046) and 1 U/ml Dispase (Stemcell Technologies, Cat#: 15569185)) for 10 min under gentle shaking and occasional gentle pipetting at 28°C. Trypsin was quenched by diluting the dissociation solution 1:2 in 50% FBS containing PBS solution. Single cells were spun down at 200 g at room temperature for 3 min, the supernatant was aspirated and cells were resuspended in 150 µl CellCarrierB. The cells were likewise measured as triplicates (48 organoids each) resulting from independent sets of organoids for each measurement.

Following RT-DC, the utilized microfluidic chips were flushed with a fluorescein-MilliQ water solution and z-stacks of the flow channels acquired with an LSM 900 Zeiss confocal fluorescence microscope (Carl Zeiss AG). For each z-stack, the pinhole size was set to one Airy unit and a Plan-Apochromat 20x/0.8 Air M27 objective was employed. The median width of each flow channel was then calculated from the z-stack using a custom Python script and the RT-DC data corrected accounting for the width of the respective flow channel.

The analysis software Shape-Out (version 2.10.0, Zellmechanik Dresden) was then used for data analysis. All samples were gated for porosity (1.0 - 1.2) and size (65 µm^2^ -160 µm^2^). Statistical analysis based on a linear mixed model, calculation of the Young’s moduli, deformation and volume as well as preparation of the data for contour plots were all carried out using Shape-Out. Plots for the volume, deformation and Young’s modulus were created using OriginPro 2021 - Update 6 (Origin Lab Corporation).

### Formation of photocleavable DNA microbeads and quantification of DNA microbead disassembly using light

Photocleavable DNA microbeads were formed in the same way as detailed above. However, 60% of the utilized linkers contained a photocleavable moiety in the center of the DNA linker sequence (for details see Table S4). In triplicates, five photocleavable DNA microbeads per sample were analyzed to quantify the breakdown of the DNA microbeads following exposure to 405 nm light. The microbeads were chosen to be 50 µm in size and imaged using 5x digital zoom. The frame time was set to 148.95 ms and the pixel size of the acquired image to 256 x 256 px. To break down the DNA microbeads, the laser power of a 405 nm confocal laser (5 mW maximum power) was set to 10% and the microbeads were continuously irradiated for 60 s, resulting in their disassembly. Analysis of the disassembly was then performed in Fiji (NIH[80]). For this, the mean fluorescence signal across the irradiated images was acquired and the data normalized to the first frame of each video. The data was plotted using OriginPro 2021 - Update 6 (Origin Lab Corporation).

### Conjugation of Wnt-surrogate proteins to DNA linkers

WNT surrogate-Fc Fusion Protein (Wnt-surrogate; ImmunoPrecise Antibodies Ltd.; Cat#: N001, Lot: 5696, 6384, 7134) was dialysed against 25 mM HEPES and 500 mM NaCl buffer at pH = 8.2 using ZelluTrans/Roth Mini Dialyzer tubes MD300 (12 – 14 kDa, Carl Roth). Dialysis was conducted at 4°C for 36 h with hourly buffer changes during the day and a long incubation overnight to remove Tris from the buffer solution. Modification of the Wnt-surrogate with an azide-moiety was achieved using an azidobutyric-NHS ester (Lumiprobe) according to the manufacturer’s recommendations. The resulting solution was then again dialysed against 25 mM HEPES plus 500 mM NaCl buffer at pH = 8.2 in the same way as before to remove any unreacted NHS-esters. DBCO-modified DNA linker strands (PC or non-PC, see Table S4) were then added to the azide-modified Wnt-surrogate in a 1:1 ratio and left to react for 76 h, yielding a final concentration of 8 µM Wnt surrogate-modified DNA linker.

### Sodium dodecyl sulfate polyacrylamide gel electrophoresis

Sodium dodecyl sulfate polyacrylamide gel electrophoresis (SDS-PAGE) was conducted using 4 - 12 % NuPAGE gels (Thermo Fisher Scientific). Prior to SDS-PAGE, the protein samples were mixed with NuPAGE denaturing agent and loading dye (1x final concentration each) and left at 95°C for 5 min to denature. Per lane, 0.65 µM of protein were applied. The samples were then loaded onto the gel and run on ice for 45 min at 150 V using 10 µL Precision Plus Protein Standards All Blue (Bio-Rad) as ladder. The gels were stained under constant shaking at 80 rpm on an Orbital Shaker D0S-10L (NeoLab) using 50 mL Coomassie stain (BioRad) overnight and imaged using a c600 gel imager (Azure Biosystems).

### Formation of DNA microbeads with photocleavable Wnt-surrogate

30 µL of a DNA microbead suspension were pelleted using a C1008-GE myFUGE mini centrifuge (Benchmark Scientific) for 2 min. Then, 20 µL of supernatant were removed to leave 10 µL of DNA microbead pellet in the reaction tube. To achieve the incorporation of DNA linker with photocleavable Wnt-surrogate, the DNA microbead pellet was resuspended with 10 µL of photocleavable Wnt-surrogate modified DNA linker (8 µM), yielding a final concentration of 4 µM modified linker. The mixture was left to incubate overnight, after which the microbeads were washed three times using 100 µL of a 1x PBS solution to remove non-incorporated DNA linkers and proteins.

### Formation of DNA microbeads with photocleavable Wnt-modified Y-motifs

30 µL of a suspension of photocleavable DNA microbeads (60% photocleavable linker DNA microbeads) were pelleted using a C1008-GE myFUGE mini centrifuge (Benchmark Scientific) for 2 min. Then, 20 µL of supernatant were removed to leave 10 µL of a DNA microbead pellet in the reaction tube.

Separately, cholesterol-modified YA-motifs were prepared by mixing the three respective single-stranded DNA strands YA-1-chol (cholesterol-modified YA-1 strand), YA-2 and YA-3 at equimolar ratios to yield a final concentration of the resulting Y-motif of 150 µM in a 1x PBS solution. Prior to mixing, the cholesterol-modified DNA single strands were left at 65°C for 1 min to break up any aggregates of the cholesterol. The Y-motifs were then annealed as described above.

To yield cholesterol-Wnt modified DNA microbeads, the cholesterol-tagged YA-motifs were mixed 1:3 with Wnt-surrogate-modified DNA linkers (DBCO-Linker, see Table S4) and incubated together for 10 min to allow for the linkers to bind to the Y-motifs.

10 µL of this solution were then added to the previously formed pellet of photocleavable DNA microbeads and mixed thoroughly. The final concentration of the Wnt-surrogate modified DNA linker was again 4 µM. The mixture was then left to incubate overnight and the microbeads likewise washed three times using 100 µL of a 1x PBS solution to remove any excess and non-incorporated DNA Y-motifs and proteins.

### Formation of DNA microbeads with photocleavable 5-fluorescein-amidite (5-FAM)

5-fluorescein-amidite-azide (5-FAM) in DMSO (Lumiprobe, 10 mM stock solution) was mixed in a 1:1 molar ratio with photocleavable DBCO-modified DNA linker (80 µM) and incubated at 4°C for 48 h to yield photocleavable 5-FAM modified DNA linkers.

To create DNA microbeads with photocleavable 5-FAM modification, 30 µL of a suspension of DNA microbeads were pelleted using a C1008-GE myFUGE mini centrifuge (Benchmark Scientific) for 2 min. Then, 20 µL of supernatant were removed to leave 10 µL of DNA microbead pellet left in the reaction tube. Next, the modified DNA linker with photocleavable 5-FAM was added at a final concentration of 4 µM to the DNA microbead pellet and the mixture incubated overnight to ensure full take-up of the modified linkers into the microbeads. The DNA microbeads were then washed three times using 100 µL of a 1x PBS solution to remove any excess and non-incorporated 5-FAM DNA linkers.

As a control, DBCO-modified DNA linkers without a photocleavable moiety were likewise modified with 5-FAM and added to DNA microbeads in the same fashion as above.

### Quantification of the release of 5-fluorescein-amidite (5-FAM) from DNA microbeads

To quantify the release of 5-FAM from the DNA microbeads, the microbeads (n = 3) were illuminated with a 405 nm laser at 10% power (5 mW maximum power) and irradiated for 42 s until the 5-FAM signal was depleted. The frame time was set to 316.51 ms and the pixel size of the acquired image to 256 x 256 px during imaging. The mean fluorescence signal of the 5-FAM dye within the DNA microbeads was then measured using the circle tool in Fiji (NIH[80]) across all frames. All data was normalized to the mean fluorescence detected in the first frame of each video and plotted using OriginPro 2021 - Update 6 (Origin Lab Corporation). DNA microbeads harboring non-photocleavable 5-FAM modified linkers were exposed to the same conditions (n = 3 microbeads) and their fluorescence signal likewise analyzed and plotted as a control to verify whether the 5-FAM depletion is due to bleaching or release.

### Sample preparation and workflow for microindentation

24 Well Glass bottom Plates (Cellvis) were treated with oxygen plasma under vacuum (0.5 mbar final pressure, 200 W) for 3 min using a 300 Semi-auto plasma processor (PVA TePla AG). To coat, the wells were next filled with 1 mL of a 0.5 mg/mL poly-l-lysine (MW = 150000 kDa - 300000 kDa, Sigma Aldrich) solution in MilliQ water, and incubated for 60 min. The wells were then washed three times using 1x PBS and 10 mM MgCl_2_ solution. 100 µL of DNA microbeads in solution were added to a total volume of 1 mL 1x PBS and 10 mM MgCl_2_ in the wells and allowed to settle and adhere to the poly-l-lysine coated wells for 30 min before washing the wells three times using 1x PBS and 10 mM MgCl_2_ solution.

Indentation experiments were conducted on a Pavone microindenter (Optics11Life) using dynamic mechanical analysis (DMA) in displacement mode. For the single microbead measurements, a cantilever with a tip size of 3 µm and a spring constant of 0.019 Nm was used. Dynamic mechanical analysis was conducted at 0.1, 0.5, 1, 2, 5, 10 and 20 Hz frequency applying an indentation amplitude of 100 nm, 2 s relaxation time between frequencies and an initial relaxation time of 10 s.

For each sample, measurements on five separate DNA microbeads were undertaken. The samples were measured as triplicates. Calculation of the effective elastic moduli was based on Hertz-model fits of the indentation curve fitted to an indentation depth equal to 16% of the cantilever-tip diameter. Further, each sample was indented no further than 5% of the sample diameter, in order to exclude any measurement artifacts based on the underlying substrate.

The data is presented as the mean ± the standard deviation of the resulting values for the effective elastic, storage and loss moduli of the different types of DNA microbeads. Data analysis was conducted using the analysis software DataViewer (V2.5.0, Optics11Life).

### Fluorescence recovery after photobleaching

Fluorescence recovery after photobleaching (FRAP) experiments were conducted and analyzed as outlined in a previous publication[48].

### Quantification of Cy3-labeled Y-motif uptake into unlabeled DNA microbeads

Unlabeled DNA microbeads were incubated with Cy3-labeled Y-motifs. Upon addition of the labeled DNA Y-motifs, the DNA microbeads were immediately imaged for 60 min. The mean fluorescence signal within DNA microbeads (n = 5) was then measured using the circle tool in Fiji (NIH[80]) at t = 0 min and t = 60 min. All data was normalized to the mean fluorescence detected at t = 0 min and plotted using OriginPro 2021 - Update 6 (Origin Lab Corporation). After imaging, FRAP measurements were conducted on three independent DNA microbeads and analyzed as described above.

### Fish husbandry and maintenance

Medaka (*Oryzias latipes*) stocks were maintained according to the local animal welfare standards (Tierschutzgesetz §11, Abs. 1, Nr. 1, husbandry permit AZ35-9185.64/BH, line generation permit number 35–9185.81/G-145/15 Wittbrodt). Fish are kept as closed stocks in constantly recirculating systems at 28°C with a 14h light/10h dark cycle. The following medaka lines were used in this study: Cab strain as a wildtype[81] and *Atoh7::EGFP*[82].

### Generation of medaka derived retinal organoids

Medaka derived retinal organoids were generated as previously described[11] with slight modifications to the procedure. Briefly, medaka primary embryonic pluripotent cells were isolated from whole blastula-stage (6 hours post fertilization) embryos[72] and re-suspended in modified differentiation media (DMEM/F12 (Dulbecco’s Modified Eagle Medium/Nutrient Mixture F-12), Gibco^TM^ Cat#:21041025), 5% KSR (Gibco^TM^ Cat#: 10828028), 0.1 mM non-essential amino acids, 0.1 mM sodium pyruvate, 0.1 mM β-mercaptoethanol, 20 mM HEPES pH=7.4, 100 U/ml penicillin-streptomycin). The cell suspension was seeded in densities of >1000 cells/per organoid (approx. 15 cells/µl) in 100µl per well in a low-binding, U-bottom shaped 96-well plate (Nunclon Sphera U-Shaped Bottom Microplate, Thermo Fisher Scientific Cat#: 174925) and centrifuged (180 g, 3min at room temperature) to speed up cell aggregation. At day 1, aggregates were transferred to fresh differentiation media and Matrigel® (Corning, Cat#: 356230) was added to the media for 9 h to a final concentration of 2%. From day 2 onwards, retinal organoids were kept in DMEM/F12 supplemented with 10% FBS (Sigma Aldrich, Cat#: 12103C), 1x N2 supplement (Gibco, Cat#: 17502048), 1mM taurine (Sigma Aldrich, Cat#: T8691), 20 mM HEPES pH=7.4, 100 U/ml penicillin-streptomycin.

Retinal organoids were either derived from embryos of wildtype Cab strain only (Figure 2B) or mixed with blastomeres of blastula-stage embryos of Atoh7::EGFP transgenic line (outcrossed to Cab) in an 4:1 ratio.

### Retinal organoid microinjection

For microinjection, day 1 retinal organoids were washed 3 times after 9 h of Matrigel incubation, transferred onto Parafilm (Thermo Fisher Scientific, Cat#: 13-374-10) and lined up against the edge of a square coverslip (24 x 24 mm) in differentiation media. Borosilicate micropipettes (1 mm OD x 0.58 mm ID x 100 mm L; Warner Instruments, Cat#: 30-0016) were pulled on a Flaming/Brown micropipette puller P-97 (Sutter instruments Co.) with the following settings: Heat 505, Pull 25, Velocity 250, Time 10, 1 cycle. The microinjection was performed with a CellTram 4m oil microinjector (Eppendorf AG) and a standard manual micromanipulator under an epifluorescence stereomicroscope (Olympus MVX10; MV PLAPO 1x objective) to visualize Cy3 fluorescently labeled DNA microbeads during microinjection.

For UV light triggered release of the DNA microbead’s cargo or disassembly of DNA microbeads themselves in alive retinal organoids, organoids kept in 100µl differentiation media on a culture dish were exposed for 60 s at a 1 cm distance to Leica EL6000 (100% intensity; Lamp: HXP-R120W/45C VIS, power input 120W, Osram Licht AG, Munich, Germany). Analysis of the disassembly was then performed in Fiji (NIH[80]). For this, the mean fluorescence signal across a ROI of the DNA microbeads position within the images was acquired and the data normalized to the first frame of the timelapse imaged retinal organoid. Wnt-surrogate release from DNA microbeads was conducted 2 h post microinjection on day 1 of retinal organoid culture, since Wnt-surrogate mediated induction of retinal pigmented epithelium was found to be only possible on late day 1.

### Embryo microinjection

Stage 20 (1 day post fertilization) embryos[72] were dechorionated using hatching enzyme, washed and kept in 100 U/ml penicillin-streptomycin containing ERM (17 mM NaCl, 40 mM KCl, 0.27 mM CaCl_2_, 0.66 mM MgSO_4_, 17 mM HEPES). Embryos were transferred onto a 1% agarose mold[83], oriented heads down for microinjection and punctured at the vegetal pole. Microinjected embryos were re-cultured on glass ware in 100 U/ml penicillin-streptomycin containing ERM until hatchling stage (s40[72]) with daily assessment of their gross morphology by stereomicroscopy.

### Fluorescent labeling and imaging

For plasma membrane staining, organoids were incubated in CellMask^TM^ Deep Red plasma membrane stain (Thermo Fisher Scientific, Cat#: C10046; 1:1000) solved in differentiation media for 30 min at room temperature (RT), washed and fixed in 4% PFA. For live nuclei staining, organoids were incubated in DRAQ5^TM^ (Thermo Fisher Scientific, Cat#: 62251; 1:500) solved in differentiation media for 30 min at RT and subsequently washed.

For whole-mount antibody staining, organoids were fixed in 4% PFA for 3 h at RT and washed with PTW (0.05% Tween20 solved in PBS). Fixed samples were permeabilized with acetone for 15 min at -20°C, blocked with 4% sheep serum, 1% BSA and 1% DMSO in PTW for 2 h at RT. Samples were incubated with primary antibodies for 48 h at 4°C. The following primary antibodies were used: chicken anti-GFP (Thermo Fisher Scientific, Cat#: A10262; 1:300), mouse anti-HuC/D (Thermo Fisher Scientific, Cat#: A21271; 1:150) and goat anti-Otx2 (R&D systems, Cat#: AF1979; 1:150). Samples were washed five times with PTW and subsequently incubated with the respective secondary antibodies (1:500) for 24 h at 4°C. The following secondary antibodies were used: donkey anti-chicken Alexa Fluor 488 (Jackson ImmunoResearch Europe Ltd., Cat#: 703-545-155), donkey anti-mouse Alexa Fluor 647 (Jackson ImmunoResearch Europe Ltd., Cat#: 715-605-151), donkey anti-goat Alexa Fluor 594 (Thermo Fisher Scientific, Cat#: A-11058), DAPI (10 µg/ml in DMSO, Carl Roth, Cat#: 6335.1). Samples were washed five times in PTW in preparation for imaging.

Fixed organoids were either imaged with Leica TCS Sp8 confocal microscope (20x oil immersion objective) or on the multiview selective-plane illumination MuVi SPIM Multiview light-sheet microscope (Luxendo Light-sheet, Bruker Corporation).[84] Live fluorescent labeling imaging of embryos were performed on the Leica TCS Sp8 confocal microscope (20x oil immersion objective). Embryos were mounted heads down in 1% low melting point agarose (Carl Roth, Cat#: 6351.5) into a MatTek dish (Mattek, Cat#: P35G-1.5-10-C) covered with ERM + 100 U/ml penicillin-streptomycin. The gross morphology of embryos was assessed by stereomicroscopy (Nikon SMZ18).

For light sheet microscopy, organoids were mounted in 1% low melting point agarose inside a FEP tube (Karl Schupp AG) fixed on glass capillaries. Four volumes (two cameras and two rotation angles) were acquired with the 16× detection setup in 2 channels: Cy3 (DNA microbeads); Far Red (CellMask Deep Red). Images were subjected to deconvolution using a theoretical point spread function.

In Figure 2D, DNA microbead fluorescent signal (Cy3) was image subtracted from Otx2 fluorescent signal (Alexa Fluor 594) prior to image overlay due to overlapping emission spectra of the spatially distinct sources of emission.

## Acknowledgements

The authors thank Dr. Sadaf Pashapour and the Microfluidic Core Facility at the Institute of Molecular Systems Engineering and Advanced Materials (IMSEAM) funded partly by the Health + Life Science Alliance Heidelberg Mannheim. The Health + Life Science Alliance provided state funds approved by the State Parliament of Baden-Württemberg. The authors thank Sebastian Fabritz and the Mass Spectrometry Facility at the Max Planck Institute for Medical Research for performing mass spectrometry experiments. The authors further thank Miroslav Tarnawski and the Protein Expression and Characterization Facility at the Max Planck Institute for Medical Research for analysis of the functionalized proteins. C.A. was partially supported by the Structured Doctoral programme (*Strukturiertes Doktorandenprogramm zum Erwerb des Dr. med. und Dr. rer. nat.*) of Heidelberg University. T.W. thanks the German National Academic Foundation (Studienstiftung des deutschen Volkes) for financial support. This work was supported by funding from the Deutsche Forschungsgemeinschaft (DFG, German Research Foundation) under Germany’s Excellence Strategy via the Excellence Cluster 3D Matter Made to Order (EXC-2082/1 - 390761711) to J.W. and K.G. K.G. was supported by the ERC starting grant ENSYNC (No. 101076997).

## Author Contributions

C.A. and T.W. contributed equally. In particular, C.A. performed the *in vivo* and *in vitro* experiments, T.W. designed, functionalized and provided the DNA microbeads.

J.W. and K.G. conceived and designed the experiments. The manuscript was written by C.A. and T.W. and edited by J.W and K.G.

