## Supplementary Information for "DNA microbeads for spatio-temporally controlled morphogen release within organoids"

**Contains:**
**Table S1-S4**
**Figure S1-S10**

**Table S1.** Volumes of the DNA microbeads and medaka retinal organoid cells as measured via real-time deformability cytometry.

| Repetition | Mean volume [$\mu m^3$] | Standard deviation [$\mu m^3$] | Triplicate mean [$\mu m^3$] | Standard deviation [$\mu m^3$] |
|---|---|---|---|---|
| **20 µM DNA microbeads** | | | | |
| **1** | 477.9 | 180.7 | 473.7 | 303.8 |
| **2** | 485 | 178.7 | | |
| **3** | 458.1 | 166.4 | | |
| **25 µM DNA microbeads** | | | | |
| **1** | 479.1 | 180.9 | 540.4 | 354.4 |
| **2** | 566.7 | 213.4 | | |
| **3** | 575.3 | 217.5 | | |
| **30 µM DNA microbeads** | | | | |
| **1** | 582.4 | 220.7 | 553.9 | 366.2 |
| **2** | 547.9 | 208.7 | | |
| **3** | 531.3 | 204.5 | | |
| **Medaka retinal organoid cells** | | | | |
| **1** | 564.3 | 168 | 555.3 | 290 |
| **2** | 547.8 | 154.6 | | |
| **3** | 553.9 | 178.8 | | |

**Table S2.** Deformation of the DNA microbeads and medaka retinal organoid cells as measured via real-time deformability cytometry.

| Repetition | Mean deformation | Standard deviation | Triplicate mean | Standard deviation |
|---|---|---|---|---|
| **20 μM DNA microbeads** | | | | |
| 1 | 0.081 | 0.016 | 0.089 | 0.035 |
| 2 | 0.089 | 0.021 | | |
| 3 | 0.097 | 0.023 | | |
| **25 μM DNA microbeads** | | | | |
| 1 | 0.098 | 0.02 | 0.084 | 0.032 |
| 2 | 0.08 | 0.018 | | |
| 3 | 0.074 | 0.017 | | |
| **30 μM DNA microbeads** | | | | |
| 1 | 0.068 | 0.017 | 0.075 | 0.03 |
| 2 | 0.08 | 0.018 | | |
| 3 | 0.077 | 0.017 | | |
| **Medaka retinal organoid cells** | | | | |
| 1 | 0.082 | 0.027 | 0.076 | 0.047 |
| 2 | 0.071 | 0.025 | | |
| 3 | 0.075 | 0.03 | | |

**Table S3** Young's moduli of the DNA microbeads and medaka retinal organoid cells as calculated from real-time deformability cytometry data.

| Repetition | Mean Young's modulus [kPa] | Standard deviation [kPa] | Triplicate mean [kPa] | Standard deviation [kPa] |
|---|---|---|---|---|
| **20 µM DNA microbeads** | | | | |
| **1** | 0.71 | 0.11 | 0.7 | 0.19 |
| **2** | 0.72 | 0.11 | | |
| **3** | 0.68 | 0.1 | | |
| **25 µM DNA microbeads** | | | | |
| **1** | 0.78 | 0.12 | 0.79 | 0.26 |
| **2** | 0.78 | 0.13 | | |
| **3** | 0.81 | 0.12 | | |
| **30 µM DNA microbeads** | | | | |
| **1** | 0.96 | 0.15 | 0.89 | 0.26 |
| **2** | 0.87 | 0.17 | | |
| **3** | 0.87 | 0.14 | | |
| **Medaka retinal organoid cells** | | | | |
| **1** | 0.81 | 0.22 | 0.88 | 0.48 |
| **2** | 0.84 | 0.19 | | |
| **3** | 0.99 | 0.38 | | |

**Table S4.** DNA sequences used for this study. The fluorescent label cyanine 3 is abbreviated as Cy3, the photocleavable group is denoted as BMN. Cholesterol is abbreviated as Chol.

| **Name** | **DNA sequence 5' - 3'** |
|---|---|
| YA-1 | GACCAACACCAGTGAGGACGGAAGTTTGTCGTAGCATCGCACC |
| YA-1-Chol | GACCAACACCAGTGAGGACGGAAGTTTGTCGTAGCATCGCACCTTT-Chol |
| YA-2 | GACCAACACCAACCACGCCTGTCCATTACTTCCGTCCTCACTG |
| YA-3 | GACCAACACGGTGCGATGCTACGACTTTGGACAGGCGTGGTTG |
| YB-1 | CAGTGAGGACGGAAGTTTGTCGTAGCATCGCACCCGACAGGAA |
| YB-1-Cy3 | Cy3-CAGTGAGGACGGAAGTTTGTCGTAGCATCGCACCCGACAGGAA |
| YB-2 | CAACCACGCCTGTCCATTACTTCCGTCCTCACTGCGACAGGAA |
| YB-3 | GGTGCGATGCTACGACTTTGGACAGGCGTGGTTGCGACAGGAA |
| Linker | GTGTTGGTCTTCCTGTCG |
| PC-Linker | GTGTTGGTC-BMN-TTCCTGTCG |
| DBCO-Linker | DBCO-TTTGTGTTGGTCTTCCTGTCG |
| DBCO-PC-Linker | DBCO-BMN-GTGTTGGTCTTCCTGTCG |

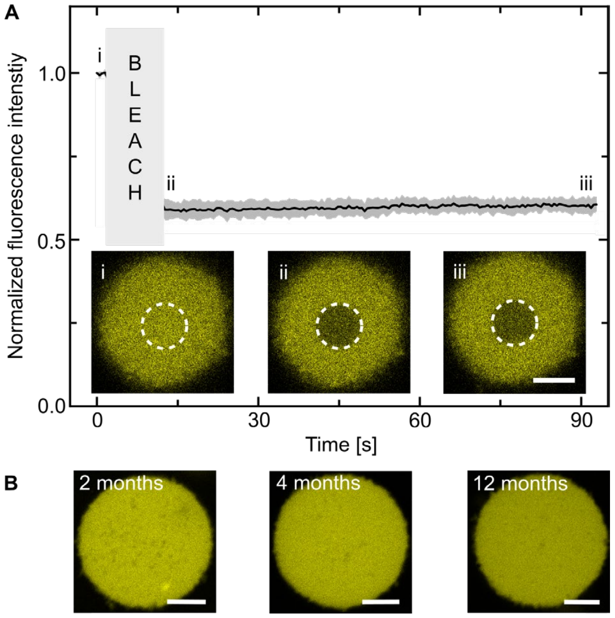


**Figure S1.** Network stability of the DNA microbeads. A) Graph showing the fluorescence recovery after photobleaching behavior of DNA microbeads (mean ± standard deviation, n = 3). Inserted confocal micrographs ($\lambda_{ex}$ = 561 nm, Cy3-labeled Y-motif, yellow) before (i) and after bleaching (ii, iii). Scale bar: 10 µm. B) Confocal microscopy images ($\lambda_{ex}$ = 561 nm, Cy3-labeled Y-motif, yellow) of DNA microbeads after 2 months, 4 months and 12 months respectively. Scale bars: 10 µm.

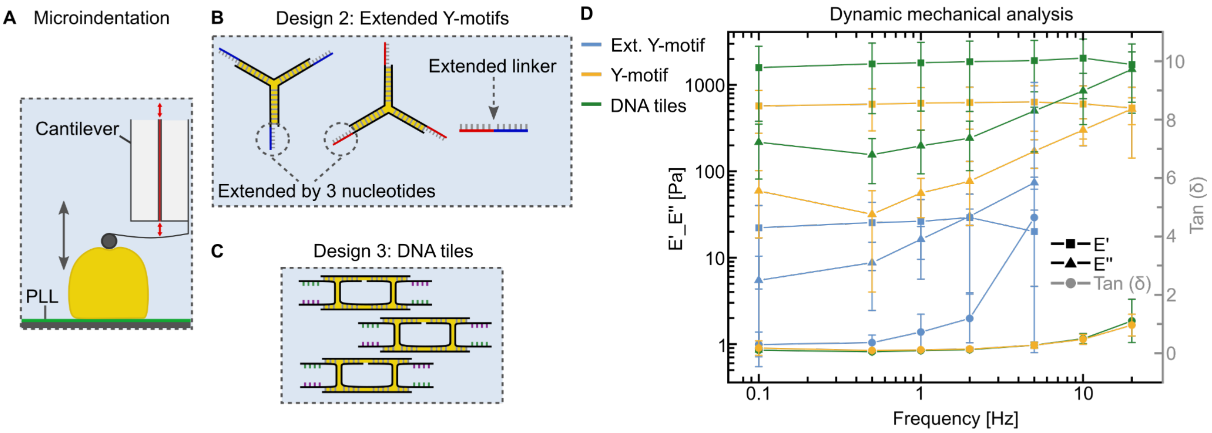


**Figure S2.** Analysis of the material properties of DNA microbeads using dynamic mechanical analysis. A) Schematic depiction of the microindentation of DNA microbeads. The DNA microbeads were attached to a poly-l-lysine-coated glass surface via charge-interaction. Indentation was then performed using a spherical cantilever tip. B) Schematic depiction of an elongated version of the Y-motif design used to form the DNA microbeads. C) Schematic depiction of a DNA tile design[85] used to form DNA microbeads via templated formation. D) Dynamic mechanical analysis of DNA microbeads with varying DNA design at 20 µM concentration across increasing indentation frequencies. The storage modulus E' (squares) and the loss modulus E" (triangles) are plotted on a logarithmic scale, while tan (δ) (E"/E', circles) is plotted on a linear scale. The values are depicted as the whole-population mean ± the corresponding standard deviation.

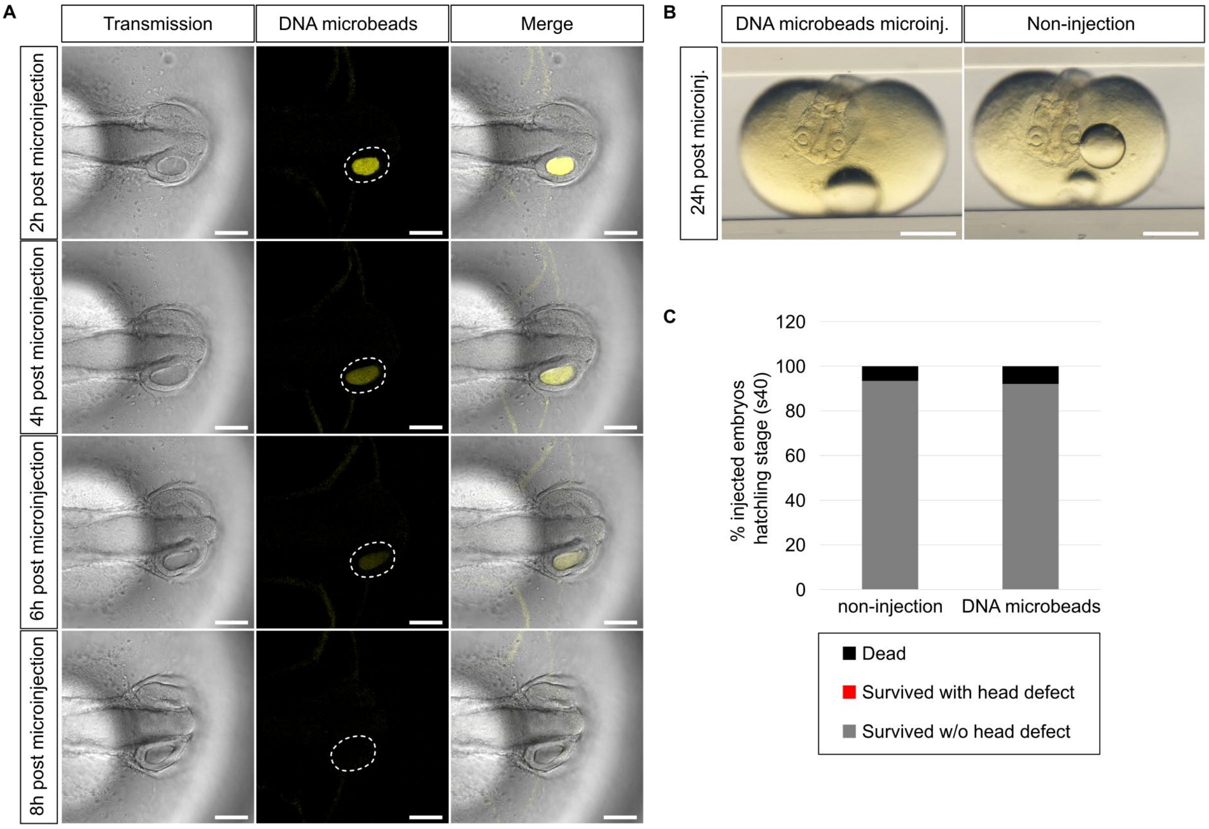


**Figure S3.** DNA microbead dynamics in the developing medaka retina. A) Representative transmission images of alive stage 20 medaka embryo 2, 4, 6 and 8 h after DNA microbead microinjection into its optic vesicle. Scale bars: 100 µm. B) Stereomicroscopic brightfield images of the embryo shown in A (DNA microbeads microinj.) and its respective uninjected control (non-injection). Scale bars: 500 µm. C) Quantification of survival and gross developmental head defects in stage 40 medaka hatchlings with (DNA microbeads; n = 25 embryos) and without (non-injection; n = 30 embryos) DNA microbead microinjection into their optic vesicles at stage 20.

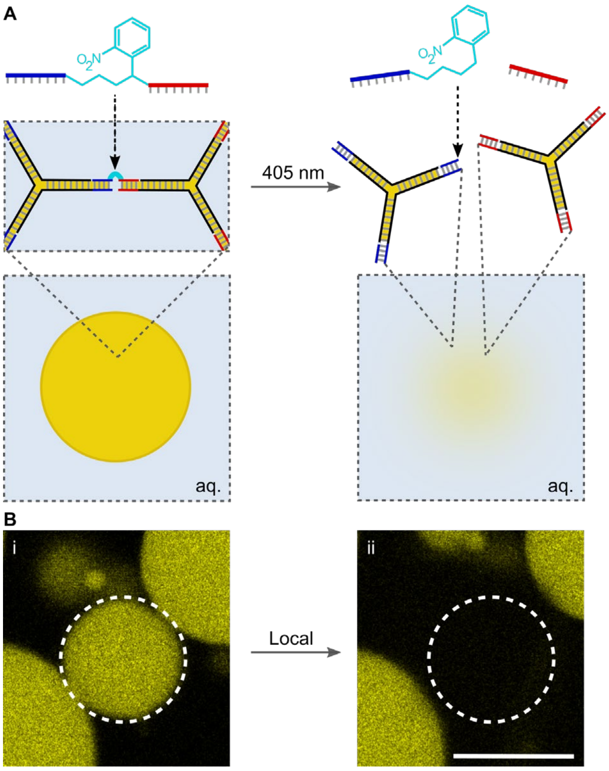


**Figure S4.** Photocleavable modification allows for locally-defined breakdown of DNA microbeads. A) Schematic illustration of DNA microbead design with an internal photocleavable group (PC) in the DNA linker sequence. B) Representative confocal images ($\lambda_{ex}$ = 561 nm, Cy3-labeled DNA) of a PC-modified DNA microbead before (i) and after (ii) irradiation with a 405 nm laser showing that only the irradiated DNA microbead was broken down by the laser. Scale bar: 20 µm.

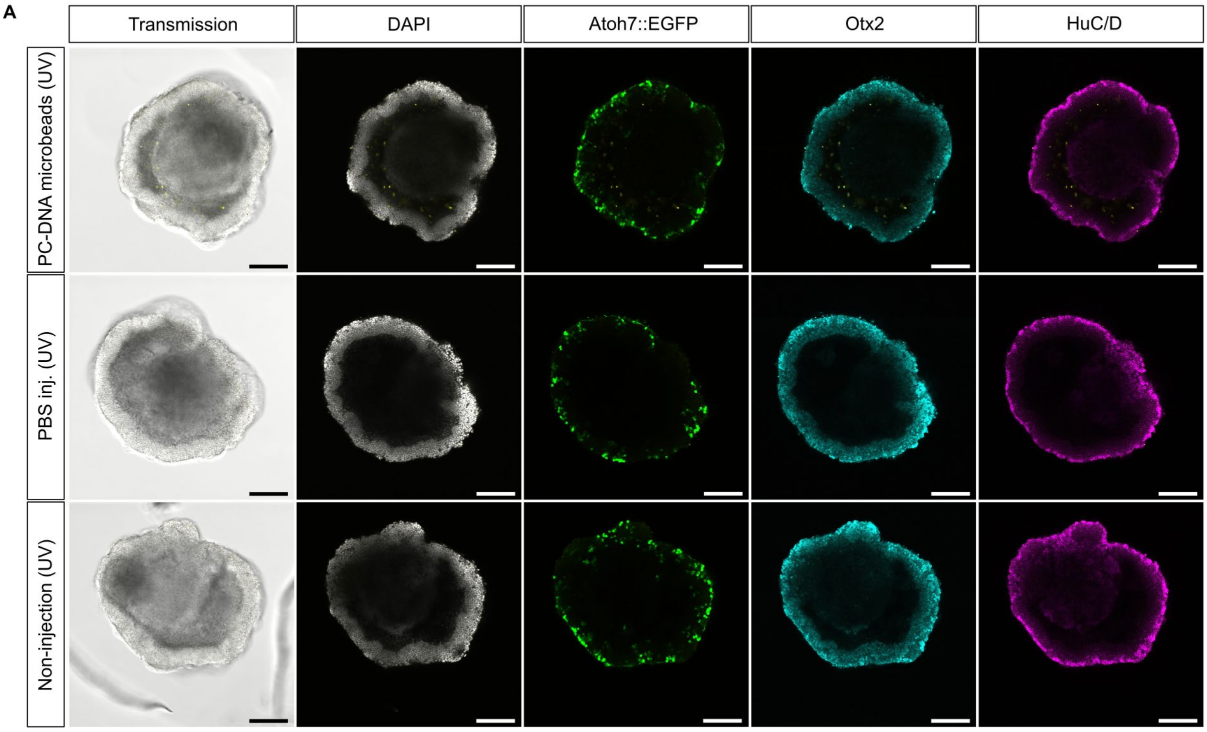


**Figure S5.** PC-modified DNA microbeads in RO development. A) Representative confocal images of whole-mount antibody stained day 4 RO (DAPI [nuclei], Atoh7::EGFP [neuroretinal ganglion cells], Otx2 [bipolar cells and photoreceptors] and HuC/D [amacrine and neuroretinal ganglion cells]) after microinjection with PC-modified DNA microbeads (PC-DNA microbeads (UV)), PBS (PBS inj. (UV)) or being left uninjected (non-injection (UV)). All conditions were exposed to the same UV light regime sufficient to trigger the DNA microbead breakdown. The microinjection and DNA microbead breakdown were conducted on late day 1 at the same timepoint used for the Wnt-surrogate release experiments in Figure 4. Representative images from n = 25 organoids across 3 independent experiments. Scale bars: 100 µm.

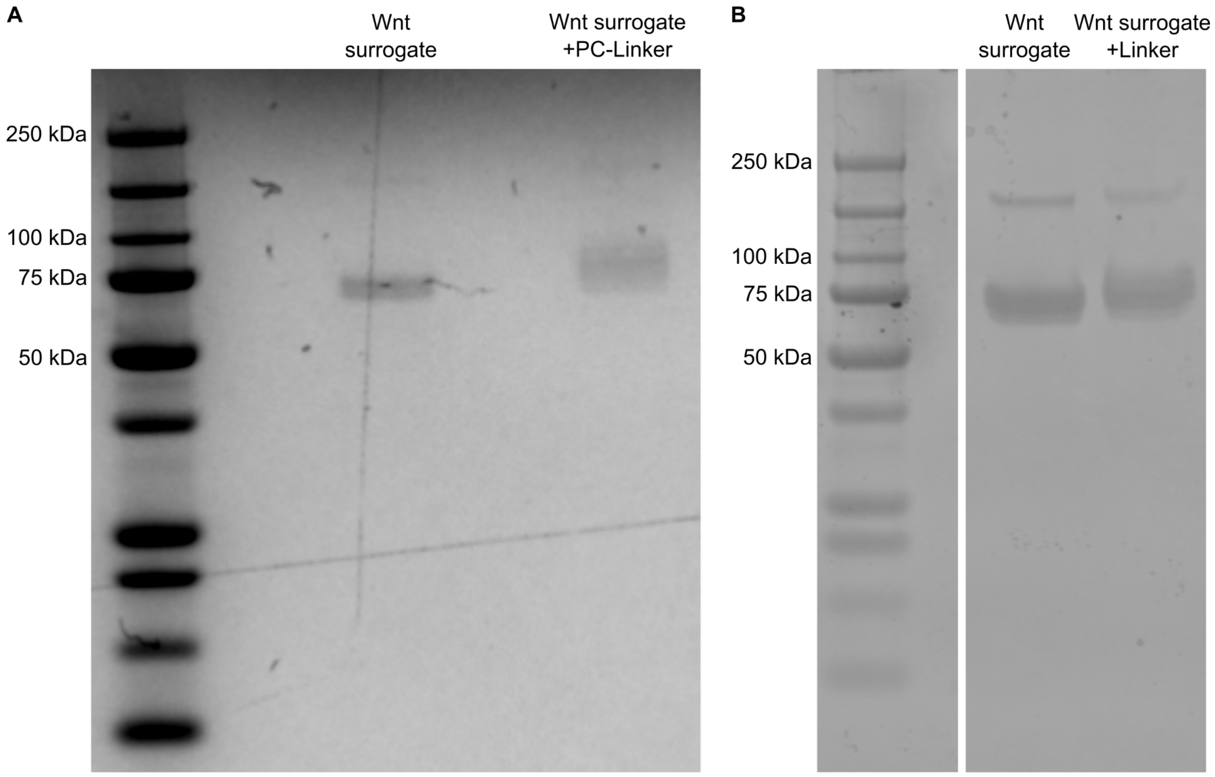


**Figure S6.** Sodium dodecyl sulfate polyacrylamide gel electrophoresis (SDS-PAGE) of the Wnt-surrogate-modified PC-Linker and DNA Linker. A) Image of an SDS-PAGE showing the non-reacted Wnt-surrogate and the PC-DNA linker-modified Wnt-surrogate. B) Image of an SDS-PAGE showing the non-reacted Wnt-surrogate and the DNA Linker-modified Wnt-surrogate.

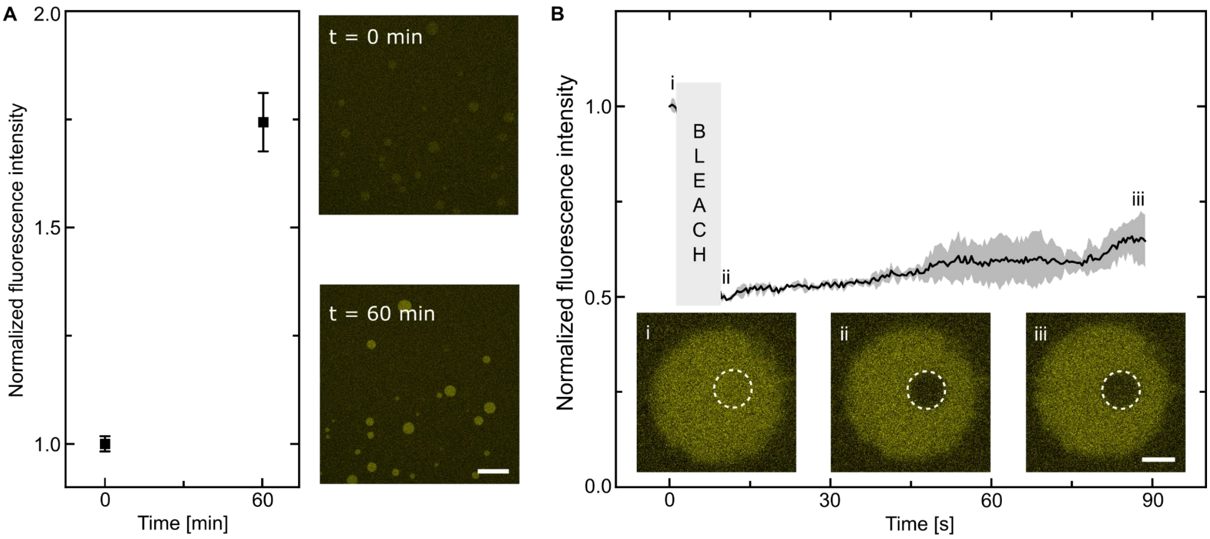


**Figure S7.** Uptake of modified DNA Y-motifs by DNA microbeads. A) Graph showing the fluorescence intensity of Cy3-modified YB-motifs upon their uptake into unlabeled DNA microbeads. Confocal microscopy images ($\lambda_{ex}$ = 561 nm, Cy3-labeled Y-motif, yellow) of DNA microbeads at t = 0 min and t = 60 min of incubation of unlabeled microbeads with labeled Y-motifs. Scale bar: 50 µm. B) Graph showing the fluorescence recovery after photobleaching behavior of originally unlabeled DNA microbeads (mean ± standard deviation, n = 3) following 1 hour of incubation with Cy3-labeled DNA Y-motifs. Inserted confocal micrographs ($\lambda_{ex}$ = 561 nm, Cy3-labeled Y-motif, yellow) before (i) and after bleaching (ii, iii). Scale bar: 5 µm.

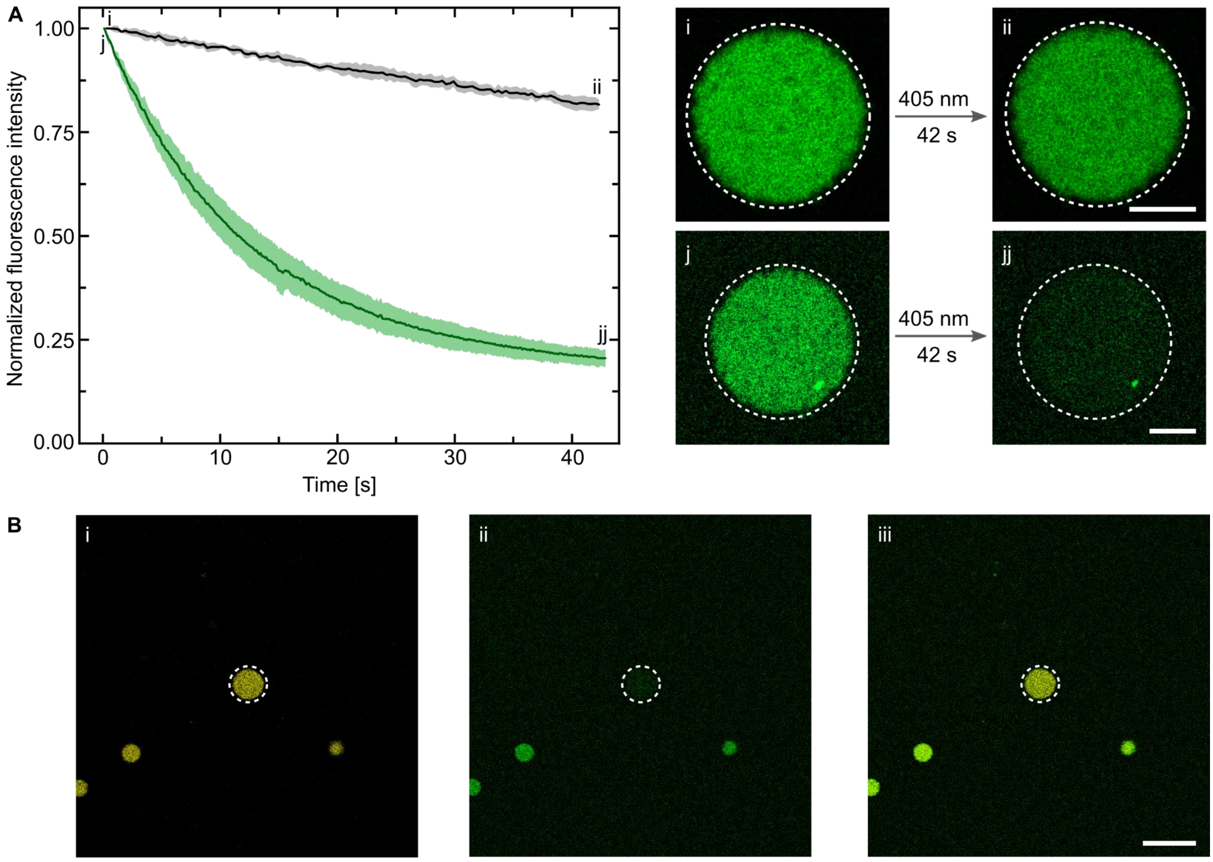


**Figure S8.** Release of 5-fluorescein-amidite (5-FAM) from intact DNA microbeads. A) Graph showing the fluorescence intensity of 5-FAM inside DNA microbeads during irradiation with a 405 nm laser for non-photocleavable 5-FAM (gray line, n = 3 microbeads) and photocleavable 5-FAM (green line, n = 3 microbeads). Insets show confocal microscopy images ($\lambda_{ex}$ = 488 nm, 5-FAM-labeled Linker, green) of non-photocleavable 5-FAM modified DNA microbeads before (i) and after (ii) irradiation with a 405 nm laser, as well as confocal microscopy images ($\lambda_{ex}$ = 488 nm, 5-FAM-labeled Linker, green) of photocleavable 5-FAM modified DNA microbeads before (j) and after (jj) irradiation with a 405 nm laser. Scale bars: 10 µm. B) Confocal microscopy images ($\lambda_{ex}$ = 561 nm, Cy3-labeled Y-motif, yellow (i), $\lambda_{ex}$ = 488 nm, 5-FAM-labeled Linker, green (ii) and overlay (iii)) of photocleavable 5-FAM modified DNA microbeads after irradiation with a 405 nm laser showing that only the irradiated DNA microbead lost its 5-FAM labeling. Scale bar: 50 µm.

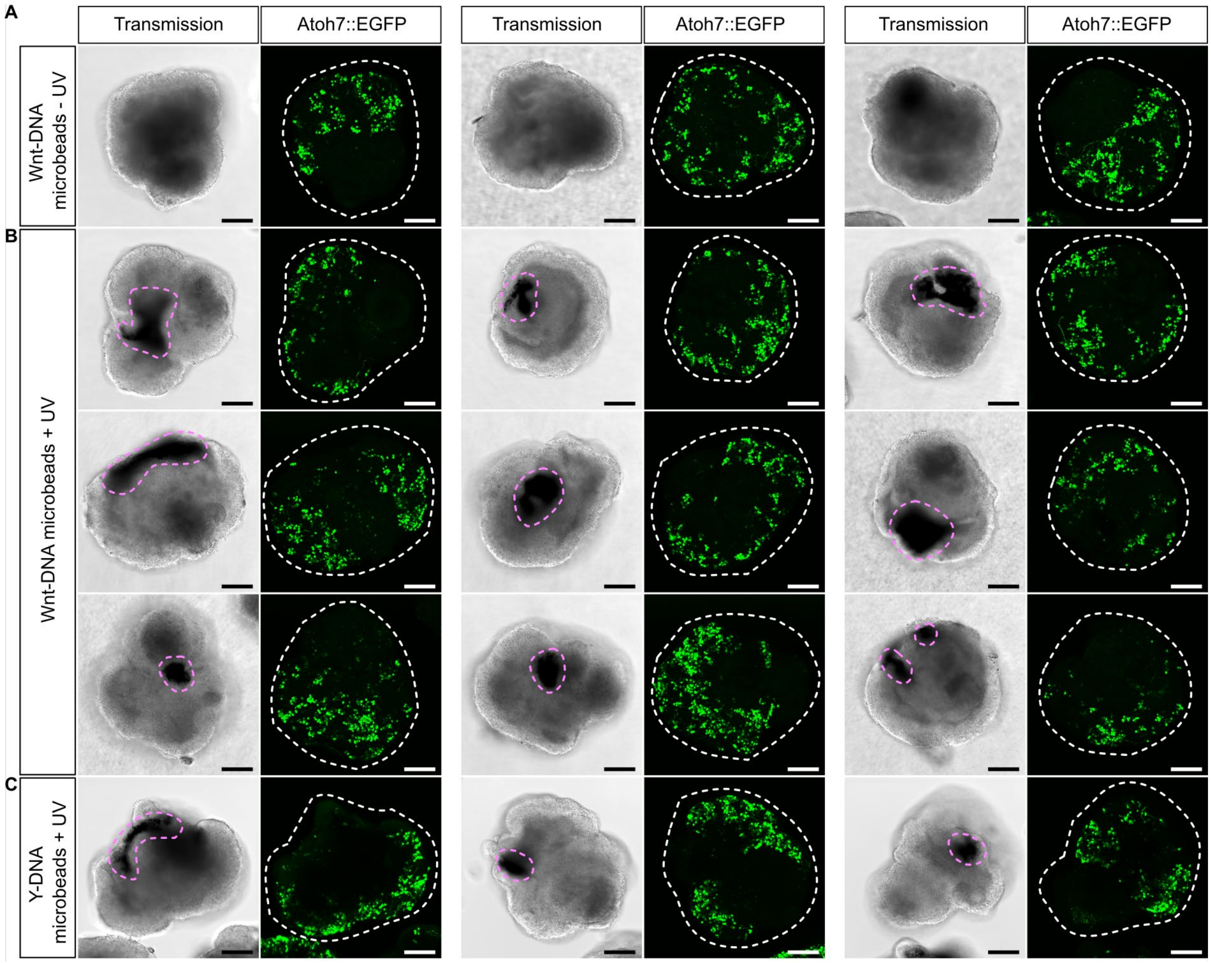


**Figure S9.** Additional representative images how the photocleavable modification of DNA microbeads permits temporally and spatially controlled morphogen release within RO. A,B) Representative confocal transmission images and maximum intensity z-projections (Atoh7::EGFP; 15 slices at 10 µm distance) of day 4 RO after Wnt-DNA microbead microinjection and Wnt-surrogate release at day 1 (Wnt-DNA microbeads + UV; DNA microbead design illustrated in Figure 4C). Respective controls (Wnt-DNA microbeads – UV) were left unexposed to UV and thus did not release their morphogen cargo. C) Representative confocal transmission images and maximum intensity z-projections (Atoh7::EGFP; 12 slices at 10 µm distance) images of day 4 RO after Y-DNA microbead microinjection and Wnt-surrogate/cholesterol-Y-DNA-motif release at day 1 (DNA microbead design illustrated in Figure 4D). Magenta dashed lines indicate retinal pigmented epithelium, while white dashed lines outline the shape of the respective RO. Representative images taken from n = 50 organoids across 3 independent experiments. Scale bars: 100 µm.

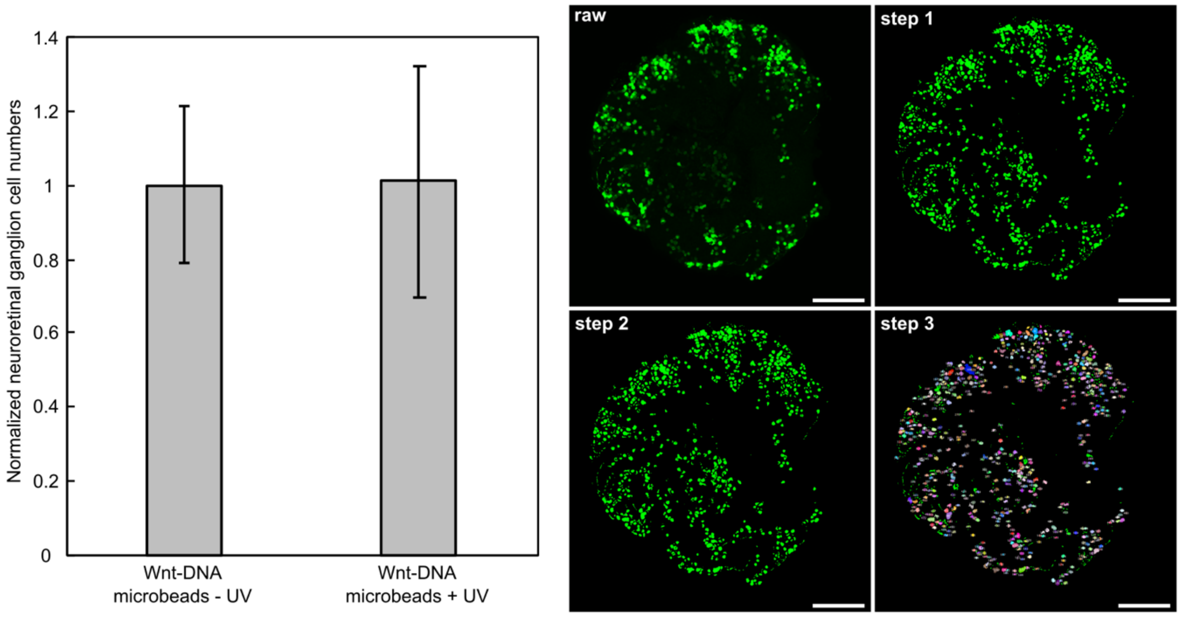


**Figure S10.** Quantification of neuroretinal ganglion cell numbers from representative images of Atoh7::EGFP maximum intensity z-projections shown in Figure 4E and Figure S9A/B. Maximum intensity z-projections of Atoh7::EGFP positive cells (Wnt-DNA microbeads - UV: n = 4; Wnt-DNA microbeads + UV: n = 10) were subjected to image processing prior to quantification[80]. Despekle noise reduction was applied to the z-projections and foreground objects were segmented from background by Auto Local Thresholding using Bernsen's thresholding with a radius of 5 and the default contrast threshold of 15 (step 1). Bernsen's local thresholding was selected due to different fluorescence intensities of cells dependent on reporter expression levels as well as non-uniform illumination dependent on cell positions within the 3D volume of the respective organoid. Thresholded binary images were Watershed to separate partially merged cell bodies by initial maximum intensity z-projections (step 2). Predominantly round cell bodies were automatically counted by Analyze Particle ImageJ implementation with a size specification of 10-infinity and a circularity specification of 0.4-1 (step 3). *Left:* Quantified cell counts were averaged and the calculated mean and standard deviation were normalized to control (Wnt-DNA microbeads - UV).
